# A Micro-Patterned, hiPSC-Derived Vascular Graft with Enhanced Endothelialization via Shear Redistribution

**DOI:** 10.64898/2026.04.12.718026

**Authors:** Jagoda Litowczenko, Yannick Richter, Piotr Paczos, Martyna Michalska, Karine Tadevosyan, Krzysztof Tadyszak, Daniel Uribe, Jose Carlos Rodriguez-Cabello, Ioannis Papakonstantinou, Angel Raya

**Author notes:** Correspondence should be addressed to: J.L. or A.R.

## Abstract

Small-diameter vascular grafts that can grow with pediatric patients and resist thrombosis remain an unmet need, primarily due to slow and unstable endothelialization. Here, we engineer a tri-layer, human induced pluripotent stem cell (hiPSC)-derived vascular graft featuring a soft, patterned lumen. We introduce a scalable soft-lithography method to imprint longitudinal micro-grooves directly into the lumen of compliant hydrogel tubes, a key advance for cell-laden constructs. Computational fluid dynamics reveals that these grooves redistribute wall shear stress into protective low-shear valleys and aligning high-shear ridges without increasing the mean load. This engineered shear landscape, combined with a bioactive elastin-like recombinamer (ELR) hydrogel matrix, synergistically enhances hiPSC-endothelial cell (hiPSC-EC) capture and retention under perfusion. Patterned grafts accelerate the formation of confluent, axially aligned endothelial monolayers with mature VE-cadherin junctions, outperforming non-patterned controls. Concurrently, smooth muscle cells within the graft wall deposit extracellular matrix, driving time-dependent mechanical maturation. This platform provides a physiologically relevant model for vascular disease and a promising strategy for engineering growth-competent pediatric grafts.

## INTRODUCTION

Cardiovascular disease remains a leading cause of mortality worldwide, and children with complex congenital lesions represent a particularly vulnerable population in need of small-diameter, growth-competent vascular conduits ^1^. Conventional synthetic grafts (e.g., ePTFE, Dacron) lack the capacity for growth and remodeling and often fail in small calibers due to compliance mismatch and thrombotic complications ^2^. Tissue-engineered vascular grafts (TEVGs) offer the promise of biological integration and somatic growth, as demonstrated by clinical studies showing neotissue formation in vivo ^3, 4^. However, a critical bottleneck limiting their success is the slow and unstable endothelialization of the luminal surface, which is essential for controlling thrombosis, inflammation, and barrier function ^5,6,7^.

Human induced pluripotent stem cells (hiPSCs) provide a transformative autologous or banked cell sources, enabling the generation of endothelial cells (ECs), smooth muscle cells (SMCs), and stromal fibroblasts to build patient-matched vascular constructs ^8^. To emulate native vessel architecture and mechanics, advanced biomaterials are required. Elastin-like recombinamers (ELRs) serve as bio-elastic building blocks that confer compliance and bioactivity ^9,10,11^, while gelatin provides a printable, integrin-adhesive base ^12,13^. Beyond biochemical ligands, physical guidance from surface topography is a potent regulator of cell behavior. Parallel micro-grooves have been shown to bias endothelial protrusion dynamics, focal-adhesion maturation, and cytoskeletal organization, promoting elongation, alignment, and retention-even under shear stress ^14,15,16^.

Despite these advances, significant gaps persist. First, robust top-down control of luminal topography in soft, fully cell-laden, and perfusable vascular constructs has not been achieved, leaving a disconnect between promising 2D contact-guidance studies and 3D graft fabrication. Second, the hemodynamic consequences and mechanistic benefits of such luminal patterning under physiologic flow remain poorly defined ^16, 17, 18, 19^. We hypothesize that integrating a precisely micro-patterned lumen into a tri-layer, hiPSC-derived, ELR-based vascular graft will create a synergistic bio-physical niche. This niche will enhance endothelial cell capture, accelerate aligned monolayer formation, and improve stability under perfusion, driven by groove-induced redistribution of wall shear stress and supported by SMC-mediated active remodeling of the graft wall.

Here we engineer a biomimetic, tri-layer vascular graft at a pediatric-relevant scale (8 mm inner diameter) from iPSC-derived cells and ELR-gelatin hydrogels. We introduce a novel soft-lithographic pipeline to imprint high-fidelity longitudinal micro-grooves directly into the lumen of hydrated, cell-laden constructs. Using computational fluid dynamics, we demonstrate how this topography redistributes shear stress to create capture-friendly valleys and alignment-promoting ridges. We then validate the performance of this integrated platform, showing that the patterned lumen synergizes with ELR hydrogel chemistry to significantly enhance hiPSC-EC retention, accelerate the establishment of confluent and junctionally mature endothelial monolayers under continuous perfusion, and promote SMC-driven mechanical maturation of the graft wall.

## MATERIALS AND METHODS

### Silicon line-pattern masters (DWL → dry etch)

Periodic line gratings were defined on Si/SiO₂ wafers by direct laser writing (DWL) (Heidelberg DWL 66+ laser writer lithography system) Wafers were solvent-cleaned, O₂-plasma activated (100 W, 60 s, O2 7 sccm), hexamethyldisilazane (HMDS)-primed, spin-coated with a photoresist (S1818 at 1.8 µm for DWL), soft-baked, then exposed via DWL to achieve 8 µm pitch pitch (4 µm groove width). After development, patterns were transferred into silicon by Bosch process in Inductively Coupled Plasma - Reactive Ion Etching (ICP-RIE) to the target heights H of ∼1.0 µm.

### Nanoimprint replication into Intermediate Polymer Stamp IPS (master → IPS)

To protect the silicon masters and enable routine replication, Intermediate Polymer Stamps (IPS; perfluoropolyether-based, Obducat-type foils) were imprinted directly from the masters by thermal/UV NIL following facility SOPs. To facilitate demoulding the masters were coated with octadecyltrichlorosilane. Imprints were cooled under load and demolded to yield flexible, wafer-scale IPS preserving pitch/height with high fidelity ^20^.

### Soft-lithographic casting of PCL daughter stamps (IPS → PCL)

Polycaprolactone (PCL, M_n ≈ 80 kDa) films were prepared as described ^21^. Briefly, PCL was dissolved at 12 wt% in chloroform (Sigma) (1 g in 5 mL; scale as needed), vortexed for 2 h at RT, cast onto glass Petri dishes and solvent-evaporated overnight. The next day, the films were gently released and thermally embossed: dishes were heated to 50 °C and IPS nanoimprint stamps (4 µm groove spacing, 1 µm height) were placed pattern-down onto PCL, then cooled to RT before carefully lifting off the IPS to obtain PCL daughter films with the groove pattern.

For forming tubular liners to cast the polyurethane acrylate (PUA) pivots, patterned PCL films were cut into rectangles sized to match the inner circumference and height of the cylindrical mold (3D-printed in transparent resin, Prusa). To obtain a PUA pivot with an outer diameter of 8.0 mm, a mold with inner diameter 8.2 mm was used to compensate the PCL thickness (∼0.10 mm per wall). Thus, each PCL sheet was cut to ∼26.3 mm (along circumference) × 20.0 mm (height), providing ∼0.5 mm overlap for a secure seam. The patterned PCL was wrapped into the mold with grooves facing the cavity. Subsequent PUA casting and curing yielded rigid PUA cylinders (“pivots”) with the pattern on the outer surface and final diameter 8.0 mm as described in the subsection below.

### Soft-lithographic casting of PUA patterned pivot (PCL → PUA)

PUA resin was formulated following a previously reported composition^22^. Briefly, diacrylate prepolymer (Ebecryl 284, Allnex) was blended with 30 wt% trimethylolpropane-ethoxylate triacrylate (Sigma 409073) as reactive diluent; photoinitiators Irgacure 184 (BASF 30472119) and 2-hydroxy-2-methylpropiophenone (Sigma 405655) were each added at 1.5 wt% relative to total resin; TEGO® Rad 2200N (1 wt%) improved wetting/release. The mixture was vortexed, degassed and stored cold, protected from light. Next the PUA was injected into the PCL-patterned tube using Positive Displacement Pipette, and UV-cured for 20 s at 365 nm (high intensity, BioX 3D Bioprinter). PCL stamps were gently demolded to yield PUA negatives with outer pattern, suitable or repeated use. Before cell work, stamps were rinsed in isopropanol, air-dried, sterilized in 70% ethanol and UV-irradiated. Patterns used in this study: we refer to patterns as F (Flat) and M1 (M4H1).

### Hydrogels topography

Bright-field images (attachment and coverage assays) were acquired on a digital microscope (Keyence VHX-series) using identical exposure settings across conditions.

### 3D printing

3D models were designed using the software Fusion 360 (Autodesk, San Francisco, California USA) for bioprinted structures as well as plastic prints. 3D printing of graft holders was conducted using the 3D printer i3 MK3S+ with Prusament PLA Galaxy Silver or a SL1S printer with Prusament Resin Tough Prusa Orange.

### Engineering of vascular grafts

Generation of the vascular constructs was performed in 3 steps. The tunica adventitia (outer layer) was 3D bioprinted with iPSC-CF, followed by casting the tunica media (middle layer) with primary aortic SMC, and the tunica intima (inner layer) seeded with iPSC-EC as schematically depicted in Figure 1.

**Figure 1.**
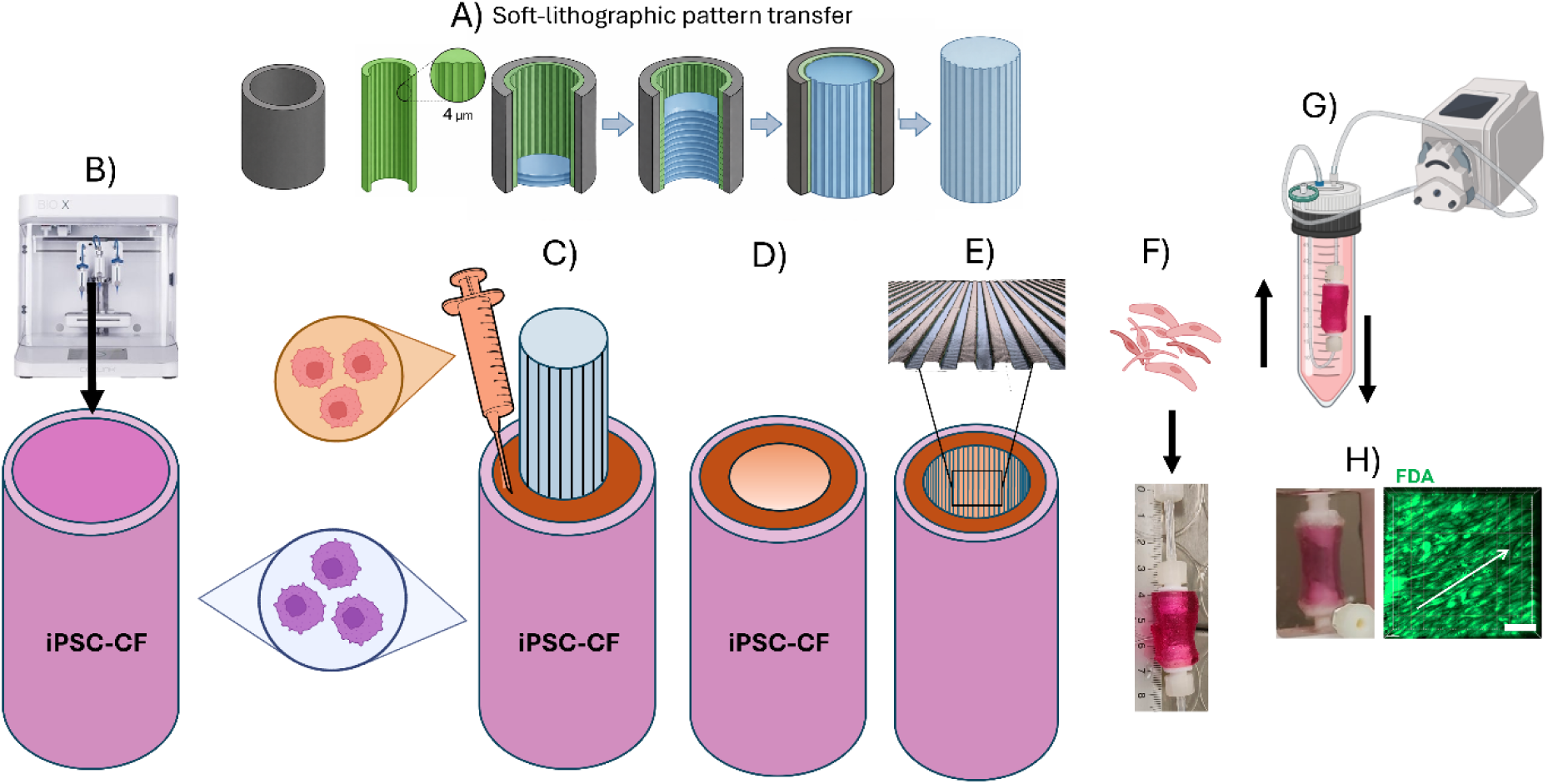
Fabrication workflow of the patterned vascular graft and perfusion set-up. (A) Soft-lithographic pattern transfer used to generate a reusable microgrooved PUA pivot via a multi-step replication chain (Direct Laser Writing → Nanoimprint Lithography → IPS → PCL → PUA), enabling high-fidelity transfer of longitudinal microgrooves (4 µm spacing, 1 µm depth). Detailed fabrication and profilometric characterization are provided in Supplementary Fig. S3. (B) Extrusion bioprinting of tubular gelatin–mTG shells containing iPSC-derived cardiac fibroblasts (iPSC-CFs), forming the outer (adventitial) layer (outer diameter ≈16 mm). (C) Casting of an ELR2-based hydrogel containing human aortic smooth muscle cells (HaSMCs) around the patterned PUA pivot to form the medial layer. (D) Removal of the pivot yields an open lumen with imprinted longitudinal microgrooves. (E) Representative microgroove topography within the lumen. (F) Seeding of iPSC-derived endothelial cells (iPSC-ECs) into the patterned lumen to establish the intimal layer. (G) Closed-loop peristaltic perfusion culture under longitudinal flow to support endothelial alignment and maturation. (H) Macroscopic view and corresponding confocal image of the luminal surface after 21 days of perfusion (FDA live staining, green), showing a confluent and aligned endothelial monolayer. Unless otherwise stated, data represent n = 4 independent experiments. Scale bar: 100 µm.

### 3D Bioprinting of the tunica adventitia

#### Bioink preparation

Gelatin from porcine skin (G1890, Sigma-Adrich) was dissolved in 60 °C preheated DMEM by vortexing and heating in a 60 °C water bath to obtain a concentration of 8% w/v. After dissolving the gelatin, the mixture was filtered with a 0.22 mm filter and left at 37 °C to keep it liquid until further usage.

In the meantime, iPSC-CF were trypsinated and, using a hemocytometer, the cell concentration determined. Cells were centrifuged at 1200 rpm for 5 min at RT, the supernatant removed, and the pellet dispersed in the gelatin solution (500.000 cells/mL). 0.15% w/v microbial transglutaminase (mTG, Ajinomoto) was added to the bioink and placed at 37°C to remove remaining micro bubbles.

#### 3D bioprinting process

The pneumatic bioprinter BIO X (Cellink) was used for bioprinting of cylindric iPSC-CF outer layer. To allow adjustment of the bioink viscosity, a temperature controlled printhead was used. The printhead was loaded with a bioink cartridge containing 3 mL bioink and a 410 μm diameter wide conical nozzle. Subsequently, tubular structures with an outer diameter of 16 mm and 2 mm thickness were printed vertically onto a specially designed printbed embedded in a 6 well plate.

As the bioink gradually solidified over time, parameters such as temperature, pressure and printing speed were adjusted during the bioprinting process to keep the bioink at a similar viscosity and enable smooth operation.

#### Fabrication of ELR-based hydrogel for tunica media

We engineered composite, imprintable hydrogels by blending gelatin with elastin-like recombinamers (ELRs) selected to decouple basal mechanics from bioactivity. DRIR (contains the slow uPA-sensitive DRIR sequence; ELR2) was used to prepare blend for casting of SMCs. These sequences and their biophysical rationale follow prior ELR design frameworks and were produced and characterized as reported previously for DRIR-bearing ELRs.

We blended gelatin with ELRs to decouple mechanics from bioactivity. ELR2 = DRIR (uPA-responsive). Production/characterization as reported previously ^10, 11^.

##### Stock solutions

Type A porcine gelatin (Sigma G1890) was dissolved in DMEM in concentration of 15% at 60°C for an hour followed by sterile-filteration (0.22 µm syringe filter). ELRs were dissolved at 50 mg mL⁻¹ in 4°C PBS for overnight (0.22 µm filtered). Phases were combined the next day 0.75 (gelatin/DMEM) : 0.25 (ELR/PBS) and cycled on ice/RT/30°C until homogenous and slightly opalescent at 30°C. ELR–gelatin hydrogels were prepared as previously described ^23^.

##### Crosslinking

Microbial transglutaminase (mTG) at final concentration of 0.15% was added to ELR-blend immediately before casting (manufacturer’s activity; details in Supplementary Methods) as previously described ^23^, to induce ε-(γ-glutamyl)-lysine crosslinking between gelatin and ELR lysine residues.

### Casting of the tunica media

For casting the tunica media, developed ELR-based hydrogel was mixed with SMC (150.000 cells/mL). Two 3D bioprinted tunica adventitia constructs were placed on top of each other and a pivot with a diameter of 8 mm inserted inside the lumen (smooth pivot or patterned-pivot. Finally, the SMC-laden ELR-based hydrogel was cast into the lumen between the 3D construct and pivot **(Figure 1).**

### Development of patterned ELR-based hydrogels

Directly after 3D bioprinting, ELR-blends were casted by positive-displacement pipette, and luers were attached. After a 3 min RT pre-set, PUA stamp was lowered without lateral sliding. Assemblies remained 10 min at RT then 5 min at 6 °C; plates returned to RT (∼40 min total) until mTG-curing completed. Stamps were peeled, gels rinsed in PBS and equilibrated in medium prior to seeding. All steps were sterile; stamps lowered vertically to avoid bubbles; only minimal finger pressure used-no external load.

### Hydrogels characterization

#### Oscillatory rheology

Small-amplitude oscillatory rheology was performed on a stress-controlled rheometer (Anton Paar MCR-302) using a 25 mm serrated parallel-plate geometry with a gap of 1.0 mm. Samples were loaded at 25 °C, trimmed, and equilibrated for 5 min. Strain-sweep tests (0.01–100%) at 1 rad s⁻¹ identified the linear viscoelastic regime (LVR). Frequency sweeps (0.1–100 rad s⁻¹) were then run at γ = 0.5% (within LVR). Temperature was controlled with a Peltier stage (±0.1 °C). Storage (G′), loss (G″) moduli and complex viscosity (η*) were exported as mean ± SD across n = 3 hydrogels per formulation.

#### CryoSEM

The morphology of all samples was analysed using an SEM-7001TTLS microscope (JEOL 204 Ltd., Akishima, Japan) operated at an accelerating voltage of 15 kV. Hydrogels (Gelatin-mTG, ELR-based hydrogel, and cell laden iPSC-CF and iPSC-SMCs) were examined using the SEM coupled with a 206 PP3000T cryo-preparation system (Quorum Technologies, Laughton, UK). This setup enabled the preparation, processing, and transfer of cryo-fixed specimens directly into the SEM chamber for observation in a vitrified state. The samples were mounted on metal holders and rapidly cryo-fixed by immersion in sub-cooled liquid nitrogen (N2 slush) at −210 °C. The frozen samples were then transferred under vacuum into the SEM preparation chamber, maintained at −180 °C. Within this chamber, the specimens were fractured to reveal fresh inner surfaces, sublimated to remove surface ice, and coated with a thin layer of platinum. Subsequently, the samples were transferred under vacuum to the SEM cryo-stage, maintained at −190 °C, where imaging of the surfaces was performed.

#### Uniaxial tensile testing

Dog-bone-like strips (width 5 mm, gauge length 15 mm, thickness 2 mm) were cut from fully cross-linked gels and tested in tension on a Zwick/Roell Z100 with a 5 kN load cell (class 1 accuracy to 0.2% Fnom). Custom flat grips used 61 HRC steel inserts coated with 95 ShA polyurethane to prevent slippage/necking of soft gels. Tests were run at 60 mm min⁻¹ from a 0.1 N pre-load to failure. True stress–strain curves were computed by engineering-to-true conversion assuming incompressibility (Poisson ∼0.5) within the large-strain regime. We report ultimate tensile strength (UTS), strain at break, Young’s modulus (tangent at 5–10% strain), and stiffness (N mm⁻¹); n = 4 per material.

### Human iPSC culture

All work with human iPSC lines was conducted under approval from the Spanish competent authorities (Commission on Guarantees concerning the Donation and Use of Human Tissues and Cells, Instituto de Salud Carlos III). The FiPS Ctrl1-mR5F-6 line (registered at the Spanish National Stem Cell Bank) was maintained on Matrigel®-coated cultureware (Corning) in mTeSR1 medium (STEMCELL Technologies). Medium was renewed daily except on the first day after passaging. For routine expansion, colonies were split 1:4–1:6 by brief incubation with 0.5 mM EDTA (Invitrogen) at 37 °C (∼2 min) and re-plated as small aggregates onto fresh Matrigel for continued maintenance.

### Differentiation of cardiac fibroblasts

Human iPSCs were differentiated into CFs in monolayer culture with sequential modulation of canonical Wnt and FGF signaling to generate second heart field progenitors as previously described ^24^. Cells maintained on Matrigel in mTeSR1 medium were dissociated into single cells with Accutase at 37°C for 8 min and seeded onto Matrigel-coated 12-well plate at a density of 1.5 million cells per well in mTeSR1 medium supplemented with 10 µM ROCK inhibitor (Y-27632). Cells were cultured in mTeSR1 medium, changed daily for 3 days. When human iPSCs achieved confluence, cells were treated with 8 µM GSK3 inhibitor (CHIR99021) in RPMI supplemented with B27 without insulin, 1% GlutaMAX, 0.5% penicillin-streptomycin, 1% nonessential amino acids, and 0.1 mM 2-mercaptoethanol (RPMI/B27-insulin medium) for 24h (day 0 to day 1). After 24 h, the medium was changed to RPMI/B27-insulin. Between day 2 and 3 the medium was changed to cardiac fibroblast differentiation basal medium (CFBM) with DMEM high glucose (4.5g/L; Corning), 500 μg/mL HSA (human serum albumin; Vitrolife), 0.6 μM linoleic acid (Sigma Aldrich), 0.6 μg/mL lecithin (Sigma Aldrich), 50 μg/mL Ascorbic Acid (Sigma Aldrich), 7.5 mM GlutaMAX (Gibco), 1.0 μg/mL Hydrocortisone Hemisuccinate (Sigma Aldrich), 5 μg/mL rhInsulin (Sigma Aldrich) supplemented with 75 ng/ml of b-FGF every other day till day 20. After, the cells were passaged in DMEM complete with 10% FBS, 1% penicillin-streptomycin and 1% GlutaMAX. iPSC-CFs were used at passages ≥ P3 for all experiments.

### Differentiation of cardiomyocytes

Human iPSCs were differentiated into CMs in monolayer culture with modulators of canonical Wnt signalling as previously described ^25^. Cells maintained on Matrigel in mTeSR1 medium (Stem Cell Technology) were dissociated into single cells with Accutase (Labclinics) at 37°C for 8 min and seeded onto Matrigel-coated 12-well plate at a density of 1.5 million cells per well in mTeSR1 medium supplemented with 10 µM ROCK inhibitor (Sigma). Cells were cultured in mTeSR1 medium, changed daily for 3 days. When human iPSCs achieved confluence, cells were treated with 8 µM GSK3 inhibitor (CHIR99021, Stemgent) in RPMI (Invitrogen) supplemented with B27 without insulin (Life Technologies), 1% GlutaMAX (Gibco), 0.5% penicillin-streptomycin (Gibco), 1% nonessential amino acids (Lonza), and 0.1 mM 2-mercaptoethanol (Gibco) (RPMI/B27-insulin medium) for 24h (day 0 to day 1). After 24h, the medium was changed to RPMI/B27-insulin and cultured for another 2 days. On day 3 of differentiation, cells were treated with 5 µM Wnt inhibitor IWP4 (Stemgent) in RPMI/B27-insulin medium and cultured without medium change for 2 days. Cells were maintained in RPMI supplemented with B27 (Life Technologies), 1% L-glutamine, 1% penicillin-streptomycin, 1% non-essential amino acids, and 0.1 mM 2-mercaptoethanol (RPMI/B27 medium) starting from day 7, with medium change every 2 days. On day 8, contracting CMs were obtained.

### Differentiation of endothelial cells

Endothelial differentiation was adapted from established lateral-mesoderm protocols ^26^. hiPSCs on Matrigel were dissociated to single cells with Accutase (37 °C, ∼8 min) and seeded onto Matrigel-coated 12-well plates at 1.5 × 10^5^ cells/well in mTeSR1 with 10 µM Y-27632 (ROCK inhibitor). After 24 h, medium was switched to EC priming medium (N2B27 composed of DMEM/F-12:Neurobasal 1:1 with N2 and B27 supplements) containing 8 µM CHIR99021 (GSK3β inhibitor) and 25 ng ml⁻¹ BMP4 (R&D Systems) for 3 days. Cultures were then transferred to EC induction medium (StemPro-34 SFM; Thermo Fisher) supplemented with 200 ng ml⁻¹ VEGF (PeproTech) and 2 µM forskolin (Sigma-Aldrich) for a further 3 days.

### Expansion and purification of endothelial cells

On day 6, cells were re-plated onto 0.1% gelatin-coated dishes and expanded in EGM-2 (Lonza) supplemented with a YAC cocktail (3 µM CHIR99021, 10 µM Y-27632, and 10 µM SB431542) for ∼4 days. Purification was performed by sequential differential detachment with TrypLE (Thermo Fisher): after two PBS rinses, cultures received TrypLE at room temperature until non-EC contaminants detached with gentle tapping and were removed; remaining ECs were then exposed to TrypLE at 37 °C (∼5 min), collected, and re-plated on 0.1% gelatin. iPSC-ECs at passages 2–3 were used for all experiments.

### Quantitative real-time polymerase chain reaction (qRT-PCR)

Total RNA was extracted from iPSC-FBs and iPSC-CMs using Maxwell® RSC simplyRNA Cells kits on a Maxwell RSC instrument (Promega). cDNA was synthesized from 1 µg RNA using Transcriptor First Strand cDNA kit (Roche). qPCR was run on a LightCycler® 480 (Roche) with GAPDH as housekeeping control. RNA from primary HUVECs served as a positive endothelial control, and human dermal fibroblast RNA as a negative control. Primer sequences are listed in **Table S1**.

### Seeding of endothelial cells inside the construct lumen to generate a tunica intima

#### Perfusion system

6 perfusion chambers consisting of 50 mL falcons were connected to the peristaltic pump (Master Flex 7521-50 Console Drive Peristaltic Pump W/ 7518-10) through Masterflex tubing system to create a closed perfusion loop system. All elements of perfusion system (tubing, Luer connectors) were autoclaved to ensure sterility.

#### Endothelial cells seeding inside vascular graft

Luers were attached to both ends of the tubular construct and secured using surgical glue. Meanwhile, iPSC-EC were collected using accutase, resuspended in EGM-2 supplemented with YAC factors, inserted in the lumen of the construct and both luer ends closed. The construct was set into a 50 mL falcon tube with the help of a specially designed structural support (Fig. S2 M-O) and incubated at a horizontal position at 37 °C in a humidified atmosphere of 5% CO2.

### Endothelial short attachment test

Tubular constructs with casted ELR-based hydrogels were used for iPSC-EC seeding. iPSC-EC (p2) were harvested and resuspended in EGM-2 in volume of 1mL. Cells were seeded inside the tubular graft at density of 150 000 cells/cm² with total number of cells 1 mln/1ml. Graft were rotated 2 times to allow iPSC-EC attachment. After 15 minutes, the flow of media was applied, and the supernatant from the wells was collected and the recovered cells counted with a hemocytometer.

### Overnight seeding and perfusion of vascular construct with endothelial cells

iPSC-EC were seeded in the lumen of the graft as previously described. After 15 minutes the falcon tube was rotated 180° to enable a more homogeneous attachment inside the lumen. The constructs were secured to the luer fittings and mounted within the perfusion chambers connected to the peristaltic pump (Master Flex 7521-50 Console Drive Peristaltic Pump W/ 7518-10) to create a closed perfusion loop system. 24 hours later the perfusion at 0.6 mL/min was applied for 3 days, after that perfusion was increased to 10 mL/min for 7, 14 and 21 days.

#### Immunostaining analysis

To visualize and examine the expression of specific markers from iPSC-CF, SMC and iPSC-EC embedded in the vascular construct, staining with primary and secondary antibody was performed. Primary antibodies utilized were polyclonal rabbit anti-VE-cadherin (IgG, 1:200, Invitrogen), monoclonal mouse anti-desmin (IgG1, 1:80, Sigma-Aldrich), monoclonal mouse anti-vimentin (IgM, 1:200, Sigma-Aldrich) and monoclonal mouse anti-α-smooth muscle actin (α-SMA, IgG2a, 1:400, Sigma-Aldrich). Secondary antibodies consisted of polyclonal donkey anti-mouse IgG (Alexa Fluor 647, 1:800, Jackson ImmunoResearch), polyclonal donkey anti-rabbit IgG (Fluorescein, 1:200, Jackson ImmunoResearch), monoclonal goat anti-mouse IgG1 (Alexa Fluor 568, 1:2000, Invitrogen) and monoclonal goat anti-mouse IgG2a (Alexa Fluor 488, 1:2000, Invitrogen). Furthermore, 4,6-diamidino-2-phenylindole (DAPI, 1:1000, Thermo Fisher), a fluorescent dye binding to DNA, was utilized to cell nuclei visualization. Additionally, Phalloidin (Texas Red™-X, 1:40, Life Technologies) was utilized to stain F-actin filaments.

Media were aspirated and the construct embedded in a 4 % paraformaldehyde (PFA, Sigma-Aldrich) solution overnight at 4 °C. PFA was removed, and the sample washed three times with PBS. Using a Leica VT1000 S Vibratome, the construct was sliced in 200 μm thick slides and left in Tris-Buffered Saline (TBS) until further use.

Upon staining, TBS was exchanged for blocking buffer consisting of TBS (Sigma-Aldrich) supplemented with 0.5% Triton-X (Sigma-Aldrich) and 6% donkey serum (Bio-Connect). The next day, blocking buffer was removed, and the sample covered with primary antibodies dissolved in TBS++ consisting of TBS supplemented with 0.1% Triton-X and 6% donkey serum. The primary antibody solution was aspirated and the sample washed with TBS for 15 minutes 4 times before applying the secondary antibodies dissolved in TBS++ under light protection. The secondary antibody solution was aspirated the next day and the construct slides washed with TBS for 15 minutes 4 times.

TBS was aspirated and the slides covered with DAPI solution (1:1000 in PBS) for one hour under light protection. Imaging was performed on scanning confocal microscope Zeiss LSM980 Airyscan2. For image processing FIJI and ImageJ were used. Primary and secondary antibodies used are listed in **Table S1.**

#### Live/dead assay

Live/dead working solution was prepared in DMEM without phenol red by mixing 50 µl propidium iodide (PI; 2 mg ml⁻¹ stock in PBS; Sigma-Aldrich) and 8 µl fluorescein diacetate (FDA; 5 mg ml⁻¹ stock in acetone; Sigma-Aldrich). Constructs were rinsed with PBS, incubated with the staining mixture for up to 5 min at RT in the dark, washed with PBS, and imaged immediately in phenol red free DMEM by on scanning confocal microscope Zeiss LSM980 Airyscan2.

### Cell morphology and spreading analysis

Confocal images (coverage assays) were acquired on Zeiss LSM980 Airyscan2 using identical exposure settings across conditions. iPSC-EC surface coverage (% area) was quantified in FIJI using background subtraction, auto-thresholding (Otsu), and binary area fraction within a fixed region of interest; four non-overlapping fields per condition were analyzed and averaged per well/biological replicate. Orientation was assessed from F-actin or phase-contrast images using OrientationJ, reporting angle histograms (0–180°) and rose plots; alignment indices were computed as described in Methods (binning and smoothing parameters as in Supplementary Information). Acquisition parameters (objective, pixel size, z-sampling) were held constant within each experiment.

### Quantification of cell orientation (OrientationJ → polar plots)

Based on phase-contrast of EC-images, local orientation was computed in Fiji using the OrientationJ plugin (structure tensor). Angle histograms (0–180°) were exported and normalized to the sum of bin counts. For visualization, Python/Matplotlib generated polar bar charts with the zero-angle at the top and the axis displayed from −90° to +90°. Plot opacity/border width were standardized across conditions.

#### Image processing and area-coverage calculation

Confocal/bright-field images were analyzed by an automated Fiji pipeline: 1.Preprocessing: grayscale import. 2.Adaptive thresholding: Gaussian method (blockSize = 11, C = 2) to segment cells under non-uniform illumination. 3.Morphology: closing then opening with a 5×5 rectangular kernel to fill intrusions and remove speckle. 5.Contour detection & hole filling: external contours were detected (cv2.RETR_EXTERNAL) and filled (thickness=FILLED). 6.Metric: area covered (%) = 100 × (cell-pixels)/(total pixels), where cell-pixels are non-zero entries of the filled mask. 7.Reporting: side-by-side panels (original/thresholded/filled) and per-image coverage were compiled into a multi-page PDF; a tab-delimited file (image name, % coverage) was saved for statistics. 8.Implementation used OpenCV, NumPy and Matplotlib; parameters were held constant within an experiment.

Area coverage (%) was analyzed per day by two-way ANOVA with factors Material Pattern (M1) and Flat/Smooth lumen (F, no pattern), including the interaction. Assumptions were evaluated on model residuals (Shapiro–Wilk; Brown–Forsythe). When needed, data were logit-transformed. Post-hoc contrasts used Šídák correction for (i) within-material comparisons (each pattern vs Flat) and (ii) within-pattern comparisons (each ELR vs Gelatin). As complementary kinetics metrics, we computed AUC(0–14 d) for %coverage and the time to 90–95% coverage (T90/T95) by linear interpolation between timepoints.

#### Orientation analysis (OrientationJ + Python) and area coverage (Python)

OrientationJ (Fiji, structure tensor), normalization to the sum of bins, polar plots in Python with 0° at top and axis −90° to +90°. Coverage pipeline: grayscale import → adaptive Gaussian thresholding (blockSize = 11, C = 2) → morphological close/open (5×5 kernel) → external-contour fill → % area = non-zero / total × 100; panels and TSV exported; code uses OpenCV, NumPy, Matplotlib.

#### CFD simulations

CFD simulations were performed in COMSOL Multiphysics v5.6 (Laminar Flow). A reduced sector of an 8-mm-diameter tube was used to resolve the micron-scale grooves, with no-slip at the wall and symmetry at the lateral boundaries. Smooth and grooved walls were compared under matched flow conditions corresponding to 10 mL·min⁻¹ in the full 8 mm-diameter lumen; in the reduced sector model, the inlet flow was scaled to the sector area to preserve the same mean wall shear stress. Wall shear stress (WSS) was quantified at representative locations on the smooth wall, in groove valleys, and on ridge crests; results are reported primarily as normalized ratios (τ/τ_smooth) to ensure robustness across rheological models. Full geometry definition, mesh settings and rheology parameters are provided in the Supplementary Information (Note S1; Figs. S4–S6; Table S2).

#### Statistical analysis

Unless specified, data are mean ± s.d.; technical fields per well were averaged before inference. All experiments were performed with n = 4 independent biological replicates unless otherwise stated. Data are presented as mean ± SD.

##### Primary (per-day) analysis

two-way ANOVA at each time point with factors Pattern (F, Pattern=M4), Type III SS; residual diagnostics (Shapiro-Wilk; Brown-Forsythe); log₁₀ transform if needed. Post-hoc: Šídák for simple effects (pattern vs flat).

##### Global analysis (report in SI)

linear mixed-effects model with fixed effects Material, Pattern, Day (3/7/14) and all interactions; random intercept for replicate/well (fields nested), Satterthwaite df; post-hoc Tukey/Šídák. Provide partial η² (ANOVA) or conditional R² (LMM) as effect sizes and adjusted CIs.

All tests in GraphPad Prism 10 or R (lme4/emmeans); α = 0.05.

## RESULTS

### Design and fabrication of a tri-layer, patterned vascular graft (Fig. 1)

To create a physiologically relevant, growth-competent vascular model, we developed an integrated pipeline for fabricating a cell-laden, tri-layer graft at a pediatric scale (8 mm inner diameter). The outer adventitial layer was formed by 3D bioprinting of a tubular shell of gelatin-mTG hydrogel laden with hiPSC-derived cardiac fibroblasts (iPSC-CFs) (Fig. 1A, Supplementary Fig. S1,S3). For the medial layer, we engineered a soft, elastic-dominant composite hydrogel by blending gelatin with an elastin-like recombinamer (ELR2), optimized for smooth muscle cell (SMC) encapsulation and casting (Fig. 1B).

A key advance of this work is the direct patterning of the graft’s luminal surface, which we achieved by developing a soft-lithographic replication chain (Direct Laser Writing → Nanoimprint Lithography → into IPS → polycaprolactone (PCL) daughter replication → polyurethane acrylate (PUA) stamp fabrication). In this study, the microgrooved pattern used throughout is referred to as M1, corresponding to parallel longitudinal grooves with 4 µm spacing and 1 µm depth. The IPS mold was first used to imprint patterned PCL films, which were then rolled into cylindrical liners and served as intermediate templates for casting reusable, rigid cylindrical PUA stamps (“pivots”) bearing parallel longitudinal microgrooves (4 µm groove width and 1 µm depth) (Supplementary Fig. S3). The ELR2-gelatin hydrogel containing primary human aortic SMCs was cast around this patterned pivot. After enzymatic crosslinking and careful pivot removal, the process yielded a freestanding tubular media with a lumen exhibiting high-fidelity longitudinal grooves, as confirmed by optical profilometry, with preserved feature dimensions across the replication chain (PUA pivot: 4.01 ± 0.12 µm spacing, 0.98 ± 0.05 µm depth; ELR2 lumen: 3.83 ± 0.22 µm spacing, ∼0.94 µm depth) (Fig. 1B–D, Supplementary Fig. S3).

The resulting tri-layer construct-comprising a bioprinted adventitia, a cell-laden ELR2-based media, and a micro-patterned lumen-was mounted in a custom perfusion chamber.

The resulting tri-layer construct, comprising a bioprinted adventitia, a cell-laden ELR2-based media, and a micro-patterned lumen, was mounted in a custom perfusion chamber (Fig. 1E-G). The complete, scalable workflow, from hiPSC differentiation to a perfusable graft, establishes a platform for studying endothelialization under flow (Fig. 1H).

### Micro-grooves redistribute wall shear stress without altering mean load

We hypothesized that the primary benefit of longitudinal luminal micro-grooves would be to engineer a more favorable hemodynamic landscape for EC adhesion and alignment, rather than to change the bulk hemodynamic load. To test this hypothesis, we performed computational fluid dynamics (CFD) simulations comparing a smooth lumen to one patterned with parallel grooves (4 µm groove spacing, 1 µm depth) under identical flow conditions (10 ml·min⁻¹,through an 8-mm-diameter lumen; smooth vs patterned compared at matched mean wall loading).

The simulations confirmed that the grooves had a negligible effect on the overall velocity and pressure fields (Fig. 2B–C). However, they dramatically altered the distribution of shear rate and WSS at the luminal wall (Fig. 2D–E). While the area-averaged WSS (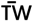) remained matched to the smooth control, the local WSS value diverged significantly depending on position relative to the groove architecture (Fig. 2D). Quantitative analysis of local wall shear stress confirmed this compartmentalization: while the mean wall shear stress remained unchanged, groove valleys experienced a pronounced reduction (τ_valley ≈ 0.25 × τ_smooth), whereas ridge crests were exposed to elevated shear (τ_ridge ≈ 1.7 × τ_smooth), yielding an approximately fivefold local shear span (Fig. 2F; Supplementary Figs. S4–S6 and Table S2), while preserving the same mean 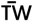 as the smooth wall. This shear compartmentalization was robust to the choice of rheological model; Newtonian and Carreau formulations produced the same ridge–valley redistribution trends (Supplementary Figs. S6–S7 and Table S1).

**Figure 2.**
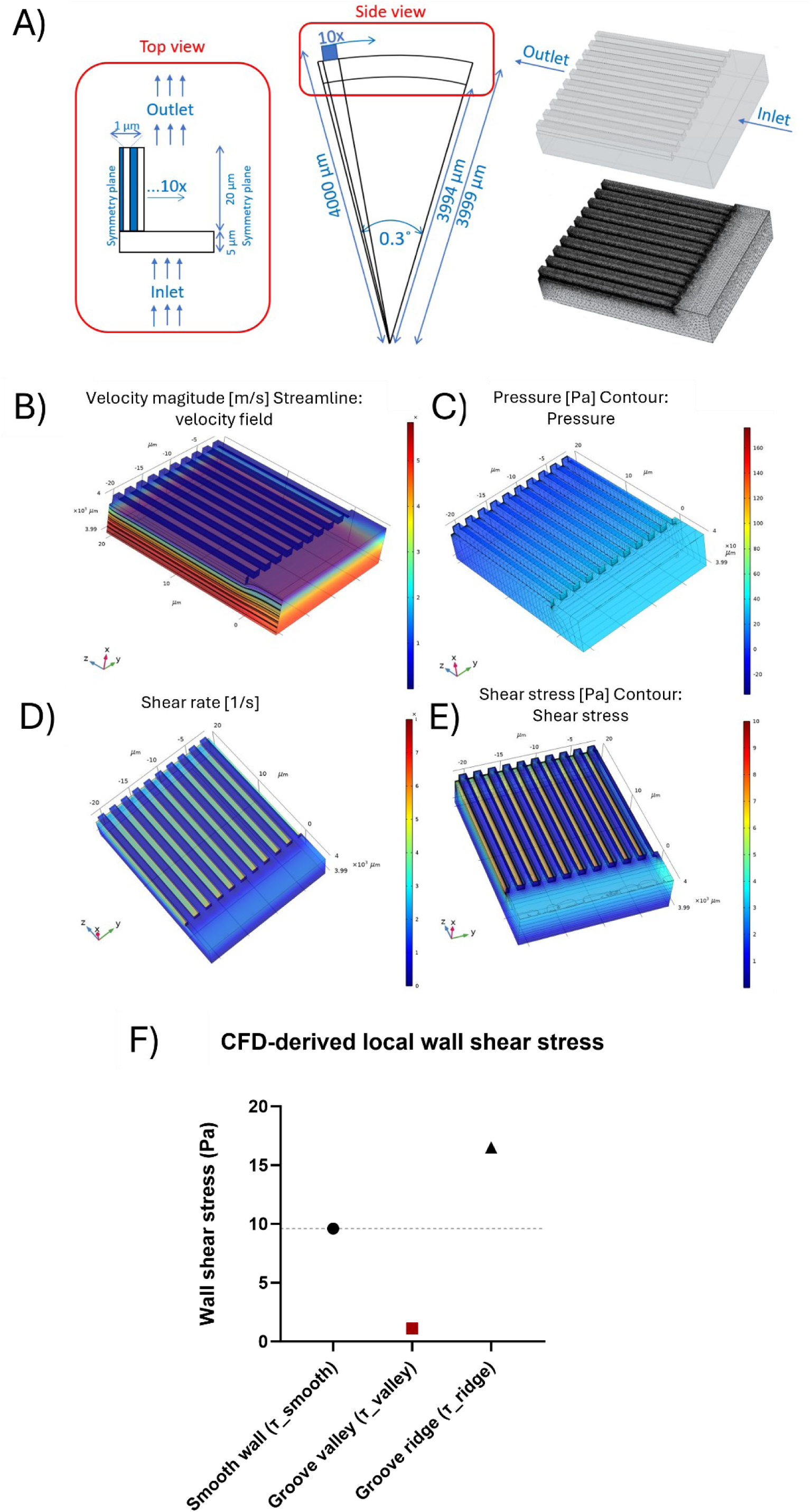
Longitudinal microgrooves redistribute wall shear stress without changing mean 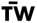. (A) Computational domain and meshing for smooth and M1-grooved lumen segments (4 µm groove spacing, 1 µm groove depth) modeled as a circumferential sector of an 8 mm-diameter lumen. Boundary conditions and symmetry planes are indicated; the grooved wall was resolved with a refined tetrahedral mesh with boundary-layer elements for accurate WSS estimation. (B) Velocity magnitude map with streamlines showing laminar, fully developed flow in both smooth and grooved cases. (C) Pressure field (Pa) demonstrating negligible difference in global pressure drop between smooth and grooved walls. (D) Shear-rate distribution (s⁻¹) showing alternating high- and low-shear regions aligned with the groove pattern. (E) Wall shear stress (WSS, Pa) contours confirming periodic high-shear ridges and low-shear valleys while preserving the same area-averaged 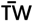 as the smooth wall. (F) Quantitative comparison of local WSS on smooth walls (τ_smooth), groove valleys (τ_valley) and groove ridges (τ_ridge), extracted from CFD simulations (n = 18 sampling points). Data are shown as box plots with individual points. Groove patterning redistributes WSS into low-shear valleys and high-shear ridges without altering mean 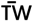.

These results provide a clear biophysical mechanism for our design: longitudinal micro-grooves redistribute wall shear stress into alternating regimes of low magnitude (valleys) and high magnitude (ridges). We posited that this engineered landscape would simultaneously offer protective niches to stabilize nascent EC adhesions during initial attachment and directional cues to promote cytoskeletal alignment along the groove (and flow) axis during subsequent maturation under perfusion ^15,16,18^.

### Validation of individual vascular graft components

With the hemodynamic rationale established, we next validated the biological and material components of the tri-layer vascular graft. We confirmed that hiPSC-derived cell types exhibited appropriate phenotypes and that the engineered ELR2-based hydrogel provided a suitable microenvironment for cell viability and function.

For the 3D-bioprinted adventitia, we differentiated hiPSC into cardiac fibroblasts (hiPSC-CFs) using sequential Wnt/FGF modulation ^24^, which yielded cells with a spindle-shaped morphology (Supplementary Fig. S1) and expression of the cardiac progenitor marker GATA4 that declined upon maturation (Fig. 3A, D), consistent with a cardiac fibroblast identity. These hiPSC-CFs were embedded in a gelatin-mTG bioink and 3D bioprinted to form the tubular adventitial layer. Crosslinker screening identified 0.15% mTG as preserving shape while maximizing viability (Supplementary Fig. S7). Live/dead staining over 1 month revealed high initial viability (94 ± 3%) and stable long-term survival (>85% after 1 month) (Fig. 3E, F). The cells progressively spread and self-organized from isolated, rounded cells into dense, interconnected multicellular networks with aligned actin cytoskeletons (Fig. 3G, Supplementary Fig. S8), demonstrating the bioink’s support of tissue-like hiPSC-CF maturation.

**Figure 3.**
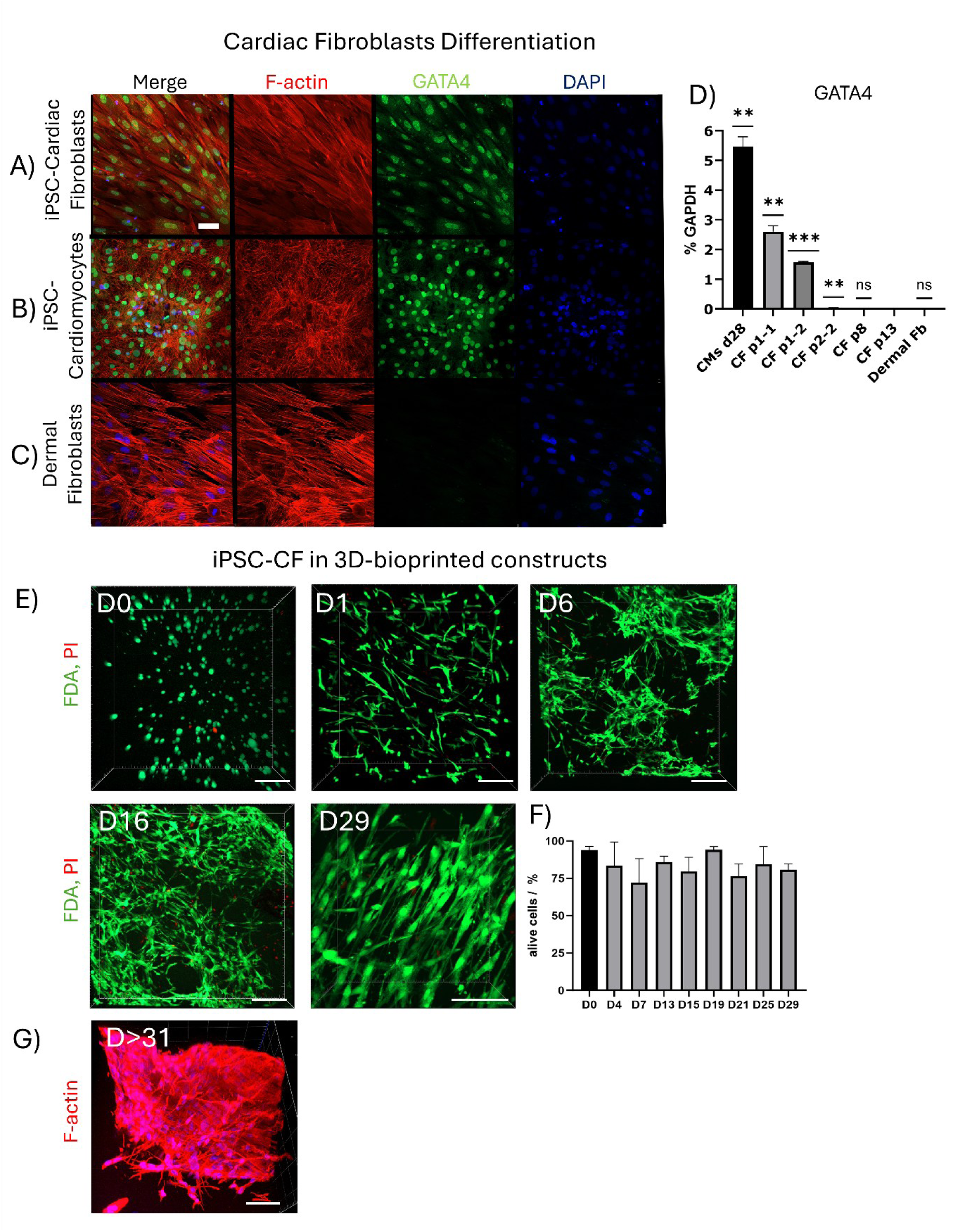
Differentiation and long-term maturation of iPSC-derived cardiac fibroblasts in 3D-bioprinted constructs. (A) Immunofluorescence staining of iPSC-derived cardiac fibroblasts (iPSC-CFs) showing F-actin (red), cardiac transcription factor GATA4 (green), and nuclei (DAPI, blue). Early-passage iPSC-CFs exhibit nuclear GATA4 expression and elongated fibroblast-like morphology. (B) iPSC-derived cardiomyocytes (iPSC-CMs) used as positive control show strong nuclear GATA4 signal. (C) Primary human dermal fibroblasts (HDFs) used as negative control lack GATA4 expression, confirming marker specificity. (D) Quantitative RT–PCR confirms elevated GATA4 transcript levels in iPSC-CFs and iPSC-CMs compared to HDFs. Data normalized to GAPDH (mean ± SD, n = 3; one-way ANOVA + Tukey). Scale bar = 50 µm. (E) 3D confocal reconstructions of iPSC-CFs within gelatin–mTG constructs over time (FDA = live cells, green; PI = dead cells, red). Cells transition from isolated spherical morphology (D0) to progressively interconnected and aligned multicellular networks (D6–D29). (F) Quantification of live cell percentage over 29 days demonstrates high initial viability (>90%) and stable long-term survival. (G) F-actin staining at ≥D31 reveals dense, highly interconnected cytoskeletal bundles consistent with advanced maturation and matrix remodeling.

The medial layer was fabricated from a composite of gelatin and the elastin-like recombinamer ELR2 (DRIR sequence), previously optimized for elasticity, and cell compatibility ^10,11^. Rheological characterization confirmed the matrix was a solid-like, shear-thinning hydrogel (G′ > G″ across 0.1–100 rad s⁻¹), ideal for casting and retaining imprinted topography (Supplementary Fig. S9A–E). Cryo-SEM revealed a fibrillar, porous microstructure distinct from gelatin alone (Fig. 4B). Primary human aortic SMCs (passage 2-3, desmin⁺/α-smooth muscle actin⁺; see Fig. 6A) encapsulated in this ELR2 hydrogel maintained high viability (95 ± 2% at day 0, 84 ± 10% by day 16) (Fig. 4F). By day 21, SMCs exhibited robust expression of contractile markers (α-smooth muscle actin, desmin) and formed extensive, interconnected bundles throughout the hydrogel volume (Fig. 4C-E, Supplementary Fig. S10), confirming a mature, functional phenotype. Depth-resolved 3D confocal imaging of the tri-layer constructs after 21 days confirmed preservation of compartmental organization across the adventitial and medial layers, with smooth muscle bundles distributed throughout the ELR2 matrix and maintaining structural continuity toward the luminal interface (Supplementary Fig. S12).

**Figure 4.**
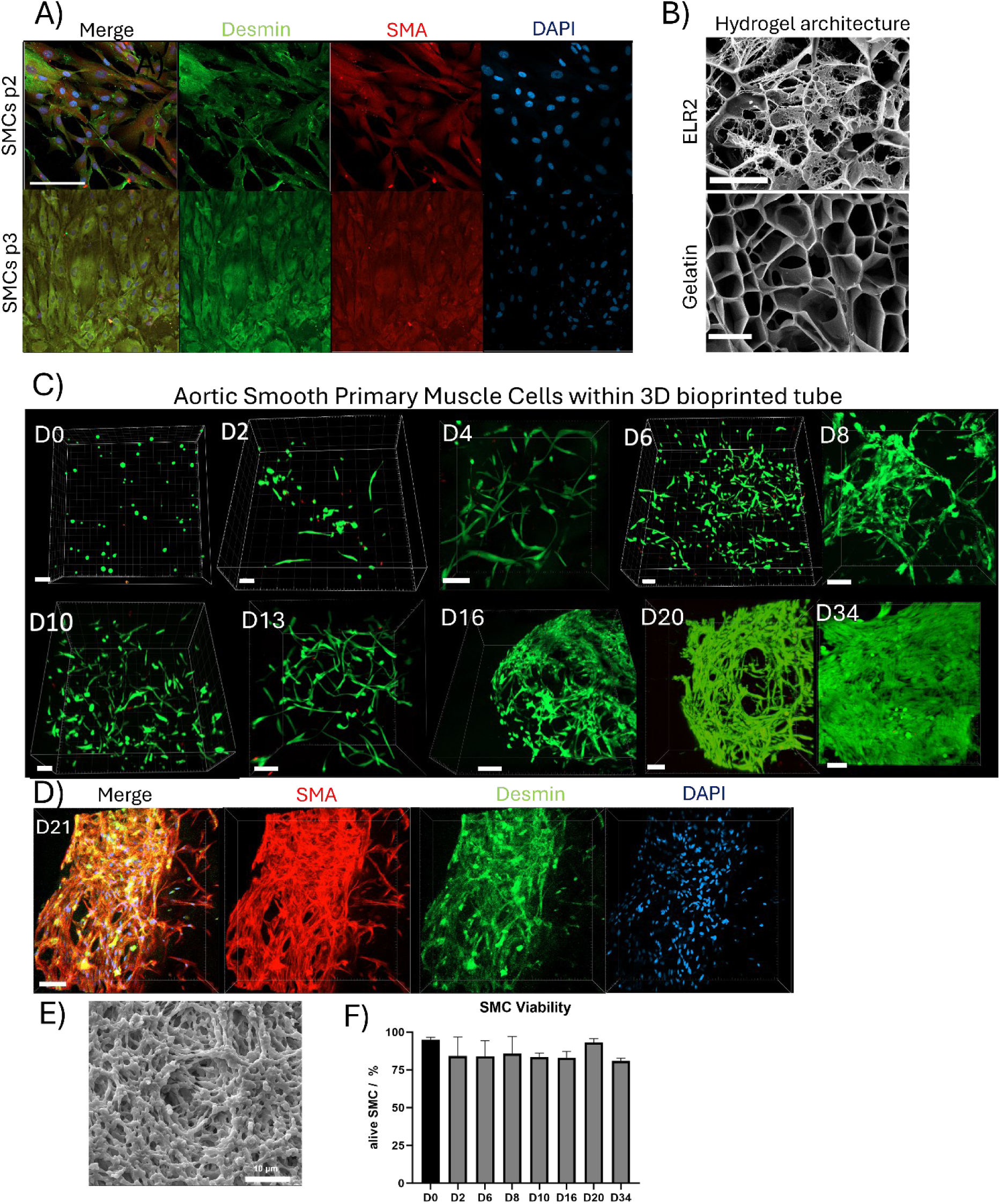
Time-dependent maturation of smooth muscle cell (SMC) layers in ELR2-based grafts. (A) Immunofluorescence for desmin (green), α-SMA (red), and DAPI (blue) confirms the contractile phenotype of primary human aortic SMCs (p2–p3). Cells display elongated morphology and well-defined actin stress fibers. Scale bar = 100 µm. (B) Cryo-SEM micrographs showing representative microstructures of crosslinked hydrogels: gelatin–mTG hydrogel with dense, irregular pore structure; ELR2/gelatin hydrogel after 24 h incubation in culture medium, showing preserved microarchitecture and pore morphology, confirming network stability after hydration. Scale bars = 10 µm. (C) Time-lapse 3D confocal reconstructions (FDA = green, PI = red) showing progressive SMC spreading, interconnection, and bundle formation from D0 to D16. Early cultures (D0–D2) show isolated cells with limited protrusions, whereas by D6–D34 extensive filamentous networks emerge throughout the hydrogel volume. Scale bar = 100 µm. (D) At D21, dense α-SMA-rich fiber bundles and desmin-positive cytoskeletal filaments dominate the construct, reflecting advanced organization and matrix remodeling under static culture. Scale bar = 100 µm. (E) Cryo-SEM micrograph showing the fibrous microarchitecture of the ELR2 hydrogel supporting cell infiltration and ECM deposition. Scale bar = 10 µm. (F) Quantification of SMC viability over time showing sustained survival (> 85 %) and stable metabolic activity.

For the luminal layer, we generated hiPSC-derived endothelial cells (hiPSC-ECs) using a Wnt/BMP-driven differentiation and purification protocol ^26^. The resulting cells displayed the characteristic cobblestone morphology of confluent endothelium and formed continuous VE-cadherin-positive junctions under perfusion (Fig. 6F). This established a reproducible and scalable source of functional ECs for lumen seeding.

### Patterned lumens enhance endothelial capture and monolayer stability under perfusion

We next tested our central hypothesis by evaluating whether the combination of ELR2 hydrogel chemistry and micro-groove topography would synergistically enhance endothelialization under physiologic shear. For this, we compared the performance of tri-layer grafts with smooth lumens against those with patterned (M1 groove) lumens under continuous perfusion following hiPSC-EC seeding.

Prior to patterned-lumen experiments, smooth hydrogel formulations were screened for early endothelial attachment and viability under static and low-flow conditions, identifying ELR2 as the optimal composition for subsequent perfusion studies (Supplementary Fig. S11).

An initial 15-minute static adhesion assay on flat surfaces confirmed the superior adhesive baseline provided by ELR2 chemistry, which retained over 90% of seeded cells, far exceeding gelatin controls (Fig. 5A). When translated to the tubular graft format under flow, the specific benefit of luminal patterning became clear. After an overnight attachment period followed by 1-hour perfusion wash (10 ml·min⁻¹), grafts with patterned ELR2 lumens exhibited near-complete cell retention (97-99% of seeded hiPSC-ECs). This represented a modest but statistically significant improvement over smooth ELR2 lumens (89–94% retention) and a marked improvement over gelatin controls (30–35%) (Fig. 5B–C). These results demonstrate that the micro-grooves enhance the early stability of the endothelial layer against shear forces, providing a robust foundation for monolayer development.

**Figure 5.**
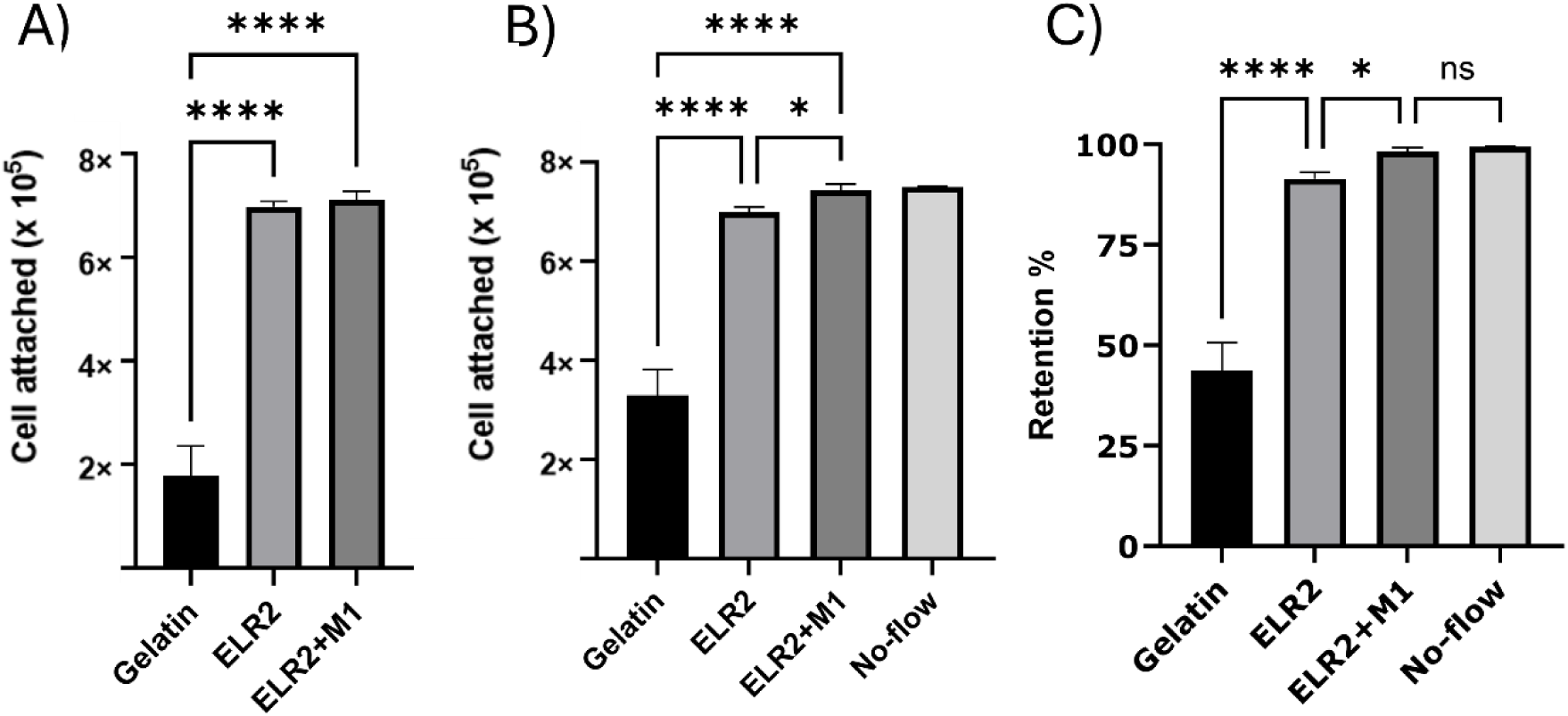
Patterned ELR2 lumens improve endothelial capture and retention under perfusion. (A) Quantification of 15-minute adhesion on flat substrates (n = 6): ELR2 ≫ gelatin; patterned ELR2 (M1) slightly higher than smooth ELR2 (one-way ANOVA + Tukey). (B) Overnight seeding followed by a 1-hour perfusion wash at 10 mL·min⁻¹: ELR2 and ELR2+M1 retained ∼0.70–0.75×10⁶ cells per tube versus ∼0.33×10⁶ on gelatin; ELR2+M1 slightly higher than smooth ELR2 (p ≈ 0.02), comparable to static control. (C) Retention relative to seeded 7.5×10⁵ cells: gelatin 35–55%, ELR2 89–94%, ELR2+M1 97–99%, no-flow ≈99% (n = 6; mean ± SD). Microgroove-induced shear gradients (Fig. 2E) explain enhanced early retention on ELR2+M1.

Monitoring endothelial behavior over 21 days of continuous perfusion revealed that this early advantage translated into accelerated and guided tissue formation. While cells on smooth lumens attached and proliferated, they displayed random orientation and slower expansion, with uncovered areas persisting for over a week. In stark contrast, hiPSC-ECs on patterned lumens rapidly and progressively aligned their morphology parallel to the groove and flow axis, achieving confluent coverage more quickly (Fig. 6A-F). Quantitative tracking of luminal surface coverage confirmed that patterned grafts achieved higher coverage at every measured time point (Days 1, 7, 14, and 21) compared to both smooth lumens and static no-flow controls (Fig. 6G-J). By the endpoint of the experiment, patterned lumens supported confluent, homogeneous monolayers that were maintained under shear, whereas areas on smooth lumens showed evidence of cell loss and monolayer thinning.

**Figure 6.**
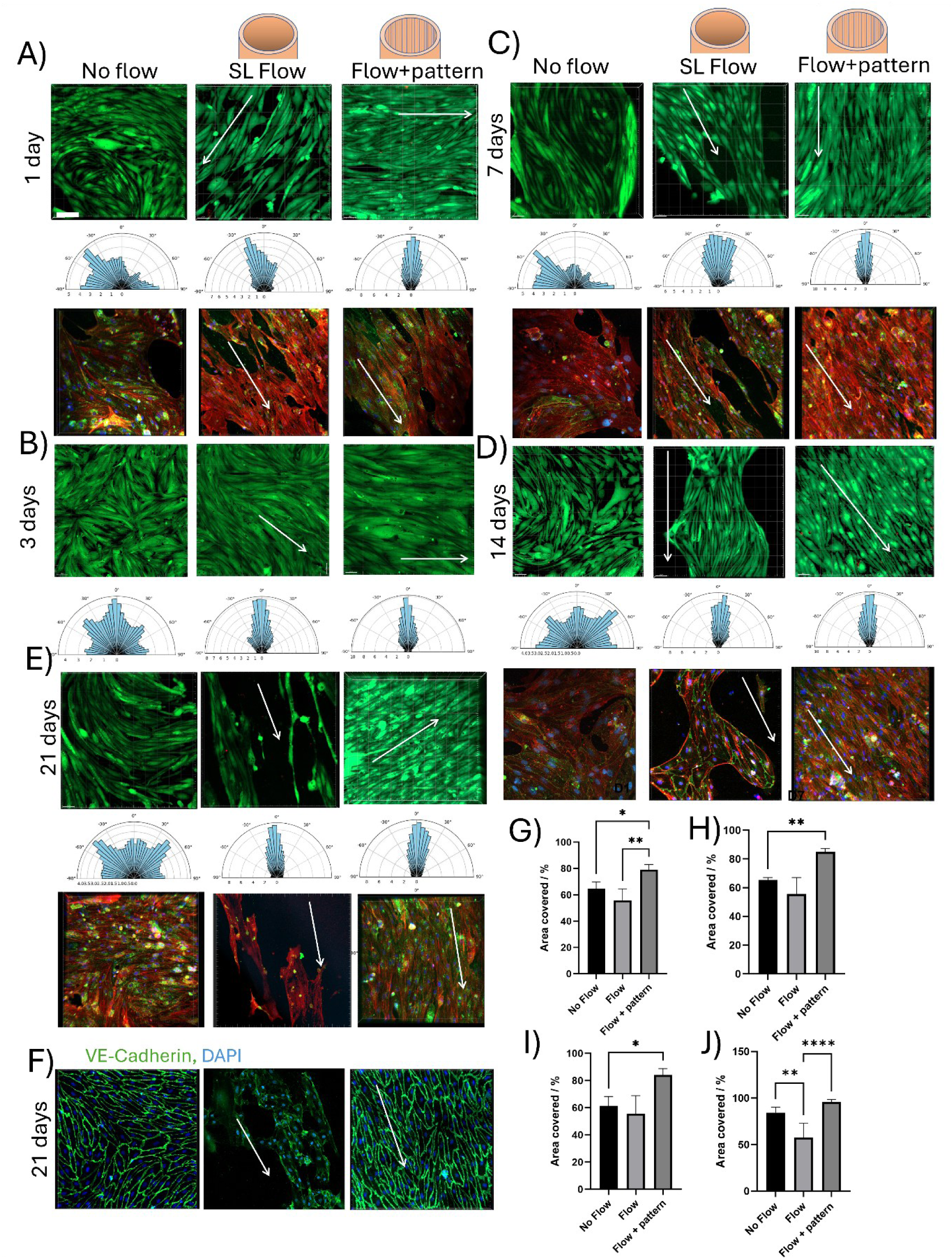
Long-term endothelialization and alignment under flow. (A–E) Time course (day 1, 3, 7, 14 and 21) of iPSC-EC monolayer formation on three lumen conditions: No flow, smooth lumen (SL) flow, and Flow + patterned lumen (longitudinal grooves, 4 µm groove spacing, 1 µm height). Top subpanels: maximum-intensity projections of the FDA stained cells illustrating cell viability and elongation (white arrow indicates dominant orientation). Bottom subpanels: OrientationJ rose plots quantifying actin-fiber angular distributions; tighter, flow-aligned distributions are evident on patterned lumens at each time point. Middle subpanels: merged immunofluorescence showing: VE-cadherin (junctional continuity) and F-actin (stress-fiber alignment).; patterned lumens exhibit earlier junction continuity and fewer uncovered regions than smooth lumens and no-flow controls. (F) Day-21 under perfusion in patterned lumen: VE-cadherin (junctional continuity). Patterned lumens display linear, continuous junctions and strongly axial actin compared with smooth and no-flow conditions. Perfusion regimen: 0.6 mL min⁻¹ for the first 3 days (conditioning), then 10 mL min⁻¹ to day 21. (G) Luminal coverage (% area) quantified over time shows patterned > smooth > no-flow at every time point (G) D1, (H) D7, (I) D14, (J) D21 (two-way ANOVA with Šídák post-hoc, p < 0.05; mean ± SD; n = 4 independent tubes per condition). Asterisks denote significance levels (p < 0.05, p < 0.01, p < 0.001). Scale bars, 100 µm.

Beyond coverage and shape, the patterned topography promoted structural maturation of the endothelium. Immunofluorescence analysis revealed that cells on patterned surfaces formed more continuous and linear VE-cadherin junctions at cell–cell borders compared to both static and smooth-lumen flow conditions, indicating the formation of stable adherens junctions (Fig. 6F). This was accompanied by a profound reorganization of the cytoskeleton; F-actin staining revealed that cells on patterned surfaces organized prominent stress fibers along the axis of alignment, while cells on smooth surfaces maintained less organized, multidirectional actin networks (Fig. 6A-E). Quantitative orientation analysis confirmed a significantly narrower angular distribution of both cell bodies and actin fibers on patterned surfaces throughout the culture period, underscoring the strength of the topographic guidance cue (Fig. 6A–E). Taken together, these results demonstrate that the micro-patterned lumen does not merely improve initial cell attachment. It actively orchestrates endothelial morphogenesis, leading to the accelerated establishment of a confluent, aligned, and junctionally mature monolayer with superior resistance to shear-induced detachment.

### Mechanical maturation of the graft wall driven by SMC-mediated active remodeling

A critical requirement for a functional vascular graft is the ability to withstand hemodynamic loads and adapt over time. We therefore investigated whether the cell-laden medial layer could actively remodel the ELR2-based matrix to enhance its mechanical properties. We performed uniaxial tensile testing on dog-bone specimens of the ELR2-gelatin hydrogel cultured with primary human aortic SMCs for 3, 7, 14, and 21 days, using acellular constructs and those seeded with non-matrix-depositing HEK293-F cells as controls. All samples exhibited J-shaped, elastomeric stress–strain curves, but key mechanical parameters evolved systematically with SMC culture time (Fig. 7).

**Figure 7.**
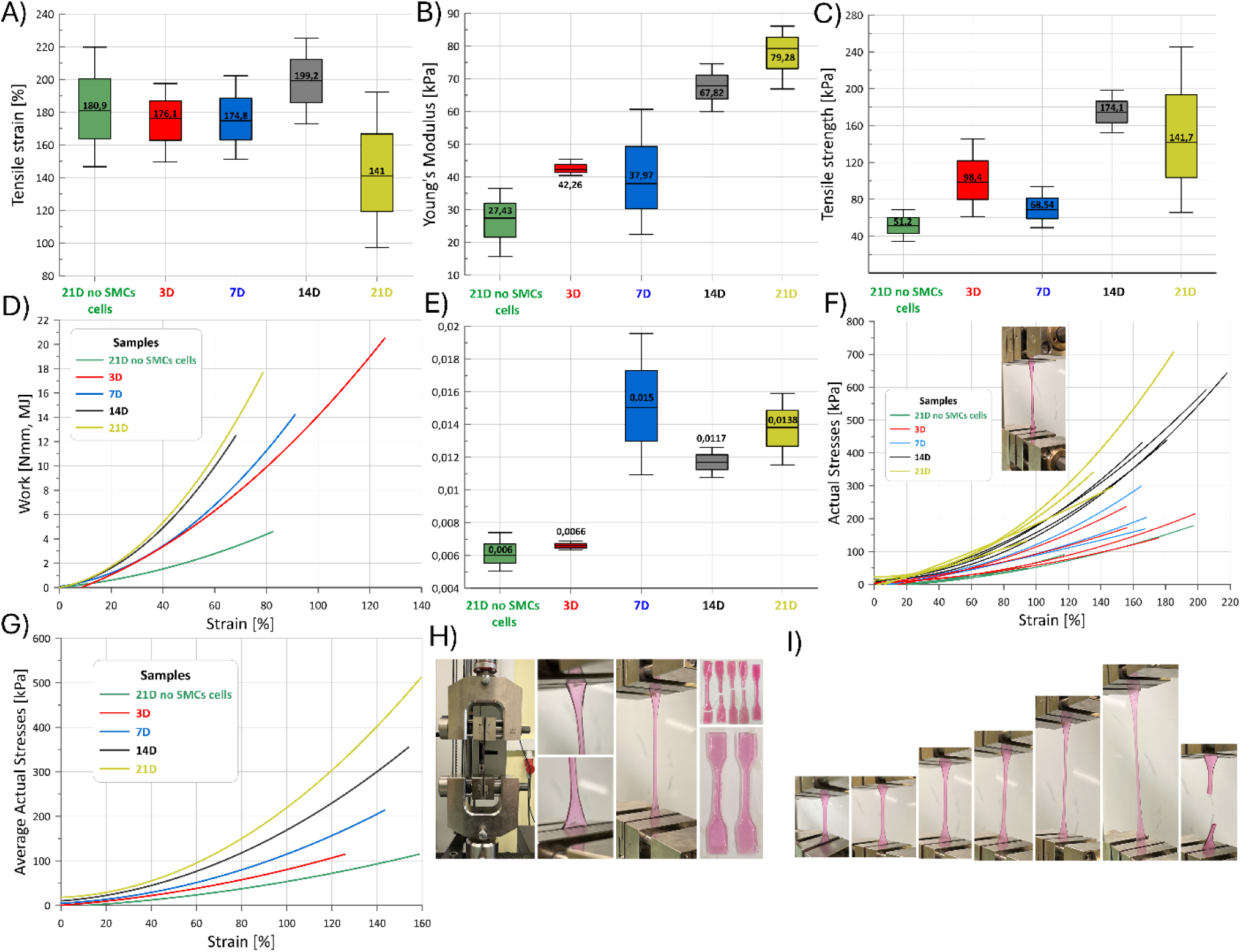
Time-dependent mechanical maturation of ELR2–SMC composites. Dog-bone specimens (35 mm total length, 5 mm width, 1.5 mm thickness) were cultured for 3–21 days under perfusion-matched conditions and tested in uniaxial tension. (A) Tensile strain at break increases up to day 14 (∼199%) before a slight decrease at day 21 (∼141%), reflecting transient matrix extensibility during remodeling. (B) Apparent Young’s modulus (small-strain region) rises progressively from ∼27 kPa (HEK293-F, 21 d control) to ∼79 kPa at day 21, indicating gradual matrix stiffening driven by SMC remodeling. (C) Ultimate tensile strength (UTS) improves significantly over time, reaching ∼174 kPa at day 14 and ∼142 kPa at day 21, surpassing the 100 kPa threshold typical for compliant vascular tissues. (D) Energy to failure (area under the stress–strain curve) shows continuous increase, confirming enhanced toughness with culture time. (E) Apparent stiffness (N·mm⁻¹) rises with SMC culture duration, peaking around day 14–21; control HEK293-F constructs remain the softest and least stiff. (F–G) Representative (F) and averaged (G) true stress–strain curves illustrate rightward and upward shifts of the constitutive response over time, consistent with progressive reinforcement of the ELR2 matrix. (H) Photographs of the mechanical testing setup and dog-bone geometry before testing. (I) Sequential frames showing deformation and failure mode characteristic of elastomeric, ductile composites.

The small-strain apparent Young’s modulus, indicative of matrix stiffness, increased progressively from ∼27 kPa in HEK293-F controls to ∼79 kPa by day 21, representing a threefold stiffening over the culture period (Fig. 7B). Concurrently, the ultimate tensile strength (UTS) rose significantly, peaking at ∼174 kPa at day 14 and maintaining a high value (∼142 kPa) at day 21, surpassing the 100 kPa threshold typical of compliant native vascular tissues (Fig. 7C).

The energy required to fracture the constructs, a measure of toughness, increased monotonically with culture duration, with 21-day SMC-laden constructs absorbing the greatest mechanical energy (Fig. 7D). In contrast, control constructs with HEK293-F cells (which are cellular but do not secrete a fibrous ECM) remained soft, weak, and brittle, with minimal change in their mechanical response over 21 days. This clear divergence confirms that the mechanical maturation is specifically driven by SMC activity and not merely a result of passive hydrogel aging or the presence of cells alone.

The temporal evolution of mechanical properties suggests a sequence of SMC-driven remodeling. An initial increase in extensibility (strain at break peaking at day 14) likely reflects early matrix reorganization and fiber engagement, followed by a phase of densification and crosslinking that enhances strength and stiffness by day 21. These data demonstrate that the ELR2 hydrogel provides a dynamic, bioactive scaffold that not only supports SMC phenotype but also facilitates cell-directed ECM assembly, leading to progressive mechanical reinforcement of the graft wall.

## DISCUSSION

Engineering a stable, aligned endothelium on soft hydrogel lumens remains a central bottleneck for vascular models and growth-competent graft concepts. Early adhesion under perfusion is fragile, and long-term organization is often lost even when initial attachment succeeds (reviewed in ^27,28,29^). Here we define a practical materials-geometry regime in which sequence-defined ELR chemistry and micron-scale longitudinal microgrooves act in concert across timescales - from minutes-scale endothelial capture to multi-week monolayer maturation under continuous flow - within an optically transparent, fully hydrated hydrogel environment. We realize this regime in a fully human, hiPSC-derived, tri-layer vascular construct fabricated by combining extrusion bioprinting of a gelatin-mTG adventitia, casting of an ELR2-based medial layer, and luminal micro-imprinting at an 8-mm inner diameter, a scale rarely achieved with living hydrogel systems.

A core mechanistic insight of our study is how luminal topography interfaces with hemodynamics. Leclech et al. (2022) demonstrated the influence of micro-grooves on endothelial cell morphology and function under flow ^16^. Our findings extend this principle into three dimensions and a dynamic graft context. CFD simulations show that longitudinal 4 µm grooves redistribute local wall shear stress without changing the mean load, generating alternating low-shear valleys (τ_valley ≈ 0.25 × τ_smooth) and high-shear ridges (τ_ridge ≈ 1.7 × τ_smooth) (Fig. 2). This “split-shear” architecture explains why patterning improves early retention even when the overall perfusion regime remains unchanged: valleys provide sheltered niches that stabilize nascent adhesions during the post-seeding phase, while ridges offer directional cues that bias cytoskeletal organization and junction alignment along the groove and flow axis. Our experimental retention and alignment results (Figs. 5-6) support this interpretation, showing that microtopography transforms a uniform shear field into a more endothelium-instructive landscape.

The early capture advantage of patterned ELR2 lumens over smooth controls under perfusion (Fig. 5) reflects this mechanism. ELR2 alone provides a strong adhesive baseline relative to gelatin, but the addition of microgrooves further improves retention under flow, consistent with geometry-enabled stabilization during initial attachment.

The choice of ELR2 as the foundational hydrogel proved critical. Fernández-Colino et al. (2019) documented the utility of ELRs as bio-elastic building blocks that impart native-like compliance and bioactivity ^9^. Our rheological characterization shows that the ELR2-gelatin composite is elastic-dominant (G′>G″) and exhibits pronounced shear-thinning behavior (Supplementary Fig. S9). These properties serve two purposes: they enable high-fidelity pattern transfer during fabrication and provide a mechanically resilient yet deformable substrate that supports cellular traction without micro-tearing. This underscores a broader design principle: viscoelasticity can be engineered to meet both fabrication and functional requirements.

The gelatin–mTG bioink used for the adventitial layer supports high initial iPSC-CF viability and long-term network formation (Fig. 3). The cast ELR2-based medial hydrogel maintains SMC viability and contractile phenotype over weeks (Fig. 4). More importantly, medial SMCs drive time-dependent mechanical maturation of the composite wall. Compared to cellular but non-remodeling controls, SMC-laden constructs show progressive increases in stiffness, strength, and toughness (Fig. 7). The Young’s modulus tripled over 21 days, and ultimate tensile strength reached ∼174 kPa, exceeding the threshold typically associated with compliant vascular tissues. This clear separation between “cells present” and “cells remodeling” confirms that mechanical evolution is biologically driven rather than a passive artifact of hydrogel aging. The construct thus matures from a printed-cast assembly into a living, adaptive tissue, a necessary step toward long-term biomechanical stability.

Translating contact guidance from flat testbeds to curved, lumen-like hydrogel interfaces at a physiologically relevant diameter required a fabrication strategy that preserves fidelity on soft, hydrated substrates while remaining compatible with tubular formats. By combining laser-written masters, replication into reusable rigid pivots, and hydrogel casting, we establish a robust route to generate high-fidelity, longitudinal microgrooves on the inner wall of a soft ELR2-based tube, validated by profilometry across the replication chain (Fig. 1; Supplementary Fig. S3). This approach integrates microfabrication with cell-compatible casting while maintaining optical transparency and mechanical compliance at 8-mm diameter.

Our findings build on established hiPSC-TEVG studies. Luo et al. (2020) showed that hiPSC-derived TEVGs can achieve mechanical strength approaching native vessels through pulsatile conditioning and SMC-mediated ECM deposition ^30^. Our platform similarly demonstrates time-dependent mechanical maturation of the ELR2-SMC medial layer, with tensile strength reaching values compatible with compliant vascular tissues. Patterson et al. (2012) highlighted the translational pathways and clinical considerations for pediatric vascular grafts, a population with a clear need for growth-competent conduits ^31,32^. Our graft, engineered at a clinically relevant pediatric scale from entirely human cell sources, directly addresses this need and provides a model for pre-clinical evaluation.

As a model, the system offers physiological diameter; human isogenic cell layers; a stable luminal endothelium under flow; and the ability to program lumen topography and medial composition. This supports disease modeling (e.g., progeria, elastinopathies) by swapping iPSC genotypes ^26,33^. For translation, iPSC sources permit HLA-typed banks or hypo-immunogenic editing to reduce rejection risk ^32,34^, and ELR-based grafts have shown promising compliance and burst metrics ^9^.

Several limitations remain. Although our perfusion regime supports stable endothelial monolayers and remodeling, physiological pulsatility and cyclic strain will likely further enhance elastogenesis and SMC matrix assembly; integrating these will be important for vascular-mechanobiology studies and translational positioning. We also have not yet evaluated hemocompatibility or inflammatory interfaces (platelet and leukocyte interactions, complement activation, and barrier function under challenge) on patterned lumens. Finally, while CFD supports the shear-redistribution principle, systematic variation of groove dimensions, alignment, and shear conditions will help refine design rules for different vessel classes and applications. Additionally, the applied flow rates are lower than those in native large vessels such as the aorta; while sufficient to establish stable endothelialization and remodeling in vitro, future studies incorporating higher, physiologically relevant flow conditions will be important for translation.

## Conclusions

We have engineered a hiPSC-derived, tri-layer vascular graft with a soft, micro-patterned luminal interface. The combination of sequence-defined ELR2 chemistry and longitudinal microgrooves establishes a hemodynamically instructive niche that guides endothelial morphogenesis from initial shear-resistant capture to confluent, aligned, junctionally mature monolayers, while the SMC-laden media undergoes progressive mechanical reinforcement. By translating contact guidance into a three-dimensional, perfusable, entirely human construct, this work provides both a versatile experimental model for vascular biology and a foundation for growth-competent pediatric vascular grafts.

## AUTHOR CONTRIBUTIONS

J.L and AR conceptualized and designed the experiments.

Y.R., M.M., K.T., P.P., D.U, H.I., JC.RC., I.P., A.R were involved in the methodology.

J.L and A.R provided resources.

J.L wrote the original draft. All authors were involved in writing, reviewing and editing.

J.L, Y.R, P.P performed statistical analysis.

J.L and A.R acquired funding and supervised the project. All authors agreed to the authorship.

## DATA AVAILABILITY

All data supporting the findings of this study are presented within the manuscript and its supplementary files. Any additional information or inquiries can be directed to the corresponding authors upon reasonable request.

## ACKNOWLEDGEMENTS

The authors are indebted to Marie Skłodowska-Curie Grant Agreement (101025242) and National Science Centre (2022/47/D/ST8/03467). PP acknowledges financial support from statutory funds of the Poznań University of Technology, granted by the Polish Ministry of Science and Higher Education (0612/SBAD/3656).

