## Supporting Information for "A Micro-Patterned, hiPSC-Derived Vascular Graft with Enhanced Endothelialization via Shear Redistribution"

**Supplementary Table 1. List of antibodies and qPCR primers used in this study.**

| Antibody | Type/Use | Host | Dilution | Manufactor |
|---|---|---|---|---|
| VE-cadherin | Primary/IF | Rabbit | 1:200 | Invitrogen, PA5-19612 |
| GATA4 | Primary/IF | Rat | 1:100 | Invitrogen, 740049T |
| Desmin IgG1 | Primary/IF | Mouse | 1:80 | Sigma, D1033 |
| SMA IgG2a | Primary/IF | Mouse | 1:400 | Sigma, A5228 |
| Phalloidin, Texas Red® | -/IF | - | 1:40 | Life technologies, T7471 |

| Primer | Sense | Antisense |
|---|---|---|
| ***GATA4*** | GGCTTACATGGCCGACGTG | CACGTCGGCCATGTAAGCC |
| ***GAPDH*** | GCACCGTCAAGGCTGAGAAC | AGGGATCTCGCTCCTGGAA |

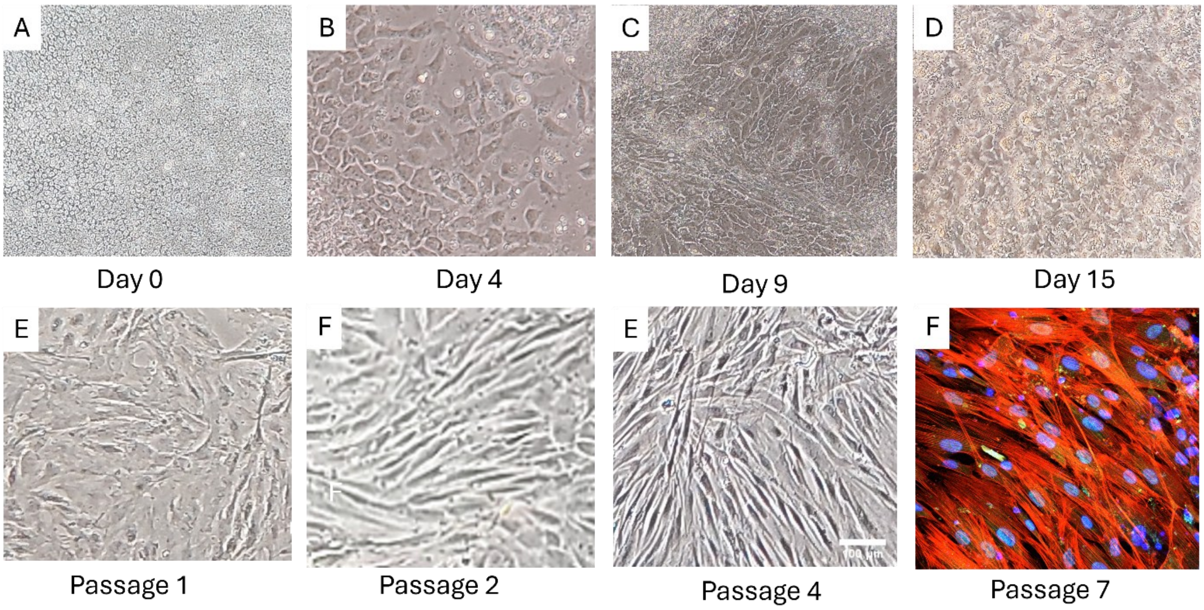


**Figure S1. Differentiation of iPSCs into cardiac fibroblasts (iPSC-CF).**
Phase-contrast images show the transition from iPSC monolayers (d0) to elongated cells (RPMI+CHIR99021, d4) and multilayer ECM-rich cultures (CFBM+bFGF, d9–15). After first passage, cells display spindle-shaped CF morphology and progressive alignment. At p7, ICC detects GATA4 (green); nuclei stained with DAPI (blue). Scale bar: 200 µm.

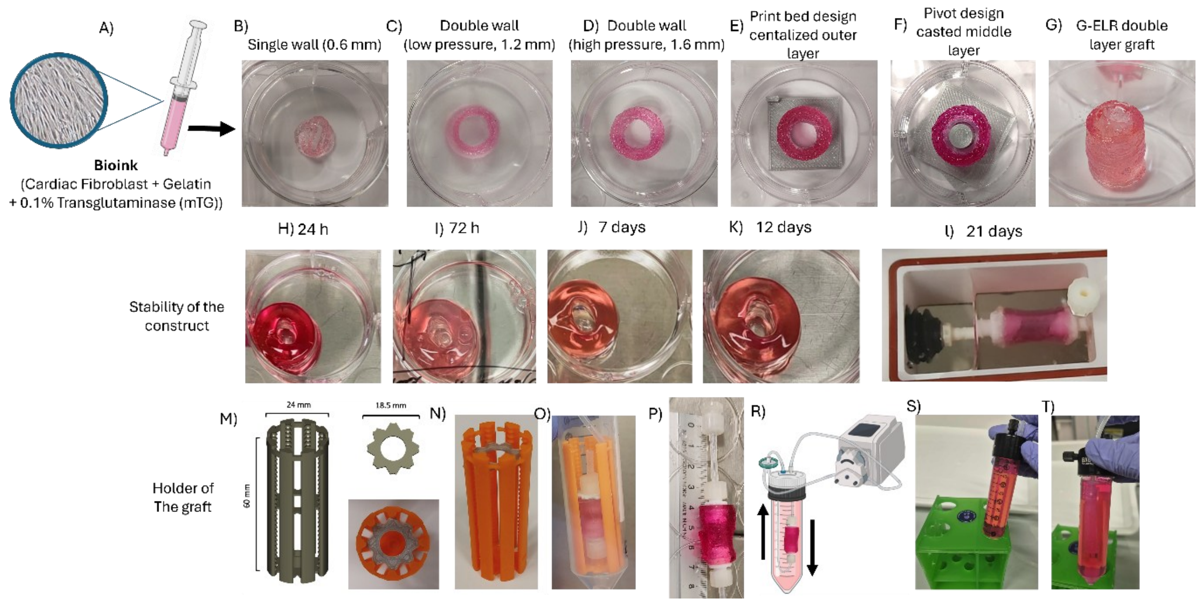


**Figure S2 | Workflow of multilayer graft fabrication and stabilization under perfusion.** (A) Schematic overview of the bioink formulation (iPSC-derived cardiac fibroblasts in 10 % gelatin + 0.1 % mTG) used to print the outer adventitial layer. (B–G) Sequential workflow of construct fabrication: (B) single-wall (0.6 mm) tubular print, (C–D) double-wall low- and high-pressure prints (1.2 mm and 1.6 mm wall thickness, respectively), (E) print-bed-centered design ensuring uniform outer deposition, (F) pivot-assisted casting of the middle ELR2 layer, and (G) final gelatin–ELR double-layer graft. (H–L) Macroscopic stability of the grafts during static incubation for 24 h, 72 h, 7, 12, and 21 days; after 21 days, constructs retained their shape and elasticity. (M–O) 3D-printed orange "graft holder" designed to secure the constructs within the perfusion chamber and prevent mechanical stress or torsion during long-term flow culture. (P) Representative graft connected to Luer fittings. (Q–R) Schematic of the closed-loop peristaltic perfusion through the graft lumen to promote endothelialization. Perfusion was applied longitudinally (top-to-bottom) to mimic aortic flow direction and minimize bubble retention. (S–T) Photographs showing the graft mounted inside the perfusion chamber (Falcon-type vessel) and connected to the flow loop.

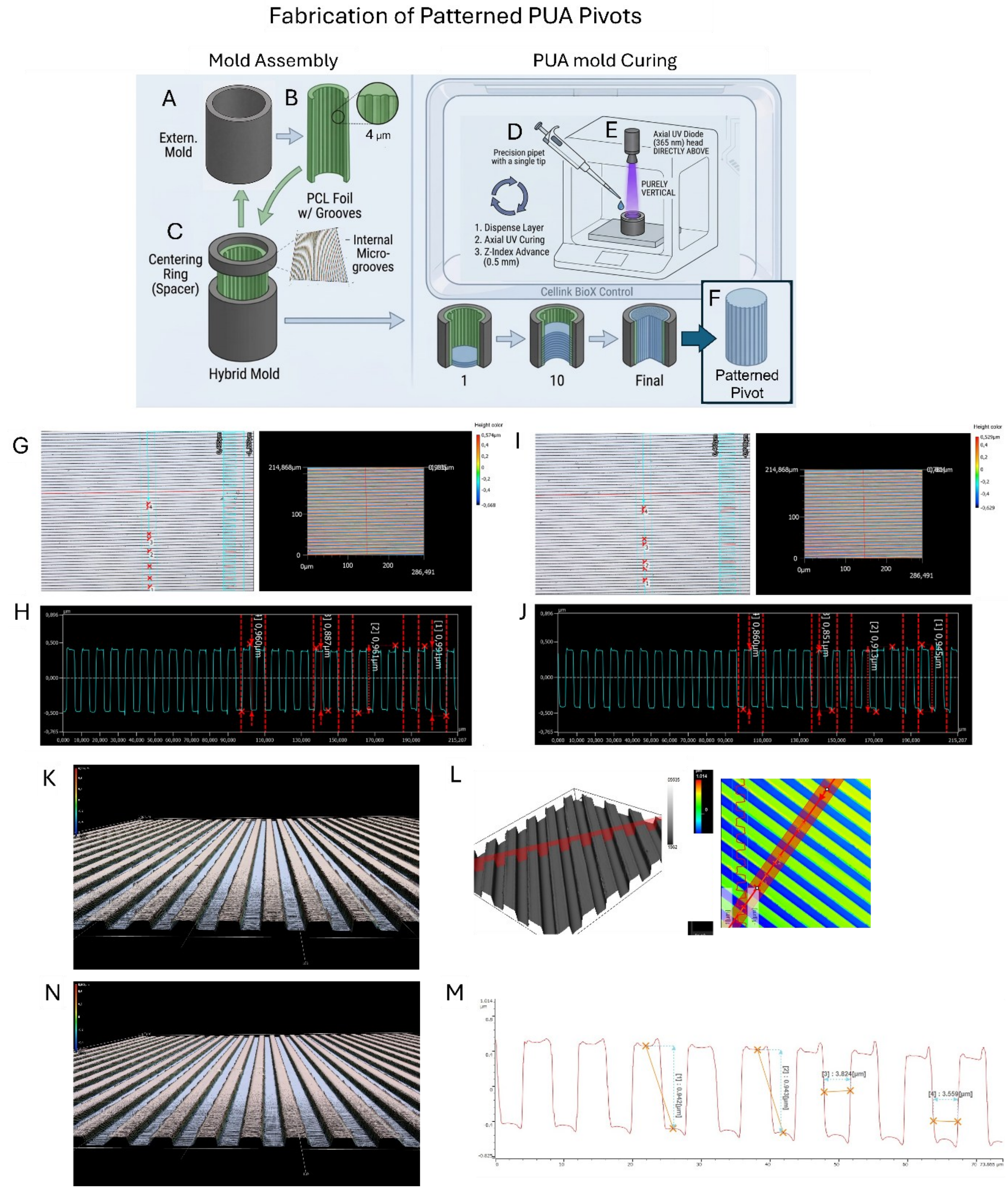


**Figure S3 | Soft-lithographic fabrication and profilometric characterization of patterned PUA pivots (PCL → PUA).**

(A–F) Schematic workflow of the soft-lithographic replication process.
(A) Cylindrical plastic mold (inner diameter 8.2 mm, height 20 mm).
(B) Patterned PCL film (4 µm pitch, ~1 µm height) prepared via nanoimprint lithography (IPS → PCL).
(C) The patterned film is cut and rolled with grooves facing inward and inserted into the mold, together with a centering ring (spacer), forming a hybrid mold.
(D) UV-curable PUA resin (Ebecryl 284 with 30 wt% TMPE-TA; 1.5 wt% Irgacure 184 and

1.5 wt% HMPP; TEGO® Rad 2200N 1 wt%) is injected into the mold cavity.
(E) The resin is crosslinked under 365 nm UV light.
(F) Demolding yields rigid PUA pivots faithfully replicating the microgrooved surface, with a final diameter of 8.0 mm.
(G–J) *Profilometric characterization of the master–replica transfer chain (Keyence VK-X3000).*
(G) 3D topography and color-coded height map of the M1 pattern embossed on PCL daughter films (4 µm pitch, ~1 µm ridge–valley amplitude).
(H) Cross-sectional profile confirming uniform periodicity and ridge alignment over >200 µm scan area.
(I–J) Corresponding topography and line profiles of the resulting PUA pivot surface demonstrate high-fidelity pattern transfer (periodicity 4.01 ± 0.12 µm, depth 0.98 ± 0.05 µm). Curvature of the cylindrical pivot was digitally compensated in Keyence software using plane-fit correction, ensuring accurate z-height reconstruction without physical sectioning.
(K–N) *Profilometry of the patterned ELR2 hydrogel lumen (Olympus DSX2000).*
(K–L) 3D height and angle maps of the hydrogel inner surface after soft-embossing from PUA pivot, showing well-defined longitudinal ridges.
(M–N) Line-profile quantification confirms faithful pattern reproduction across >200 µm field, matching the original M1 geometry (ridge-to-valley spacing ≈ 4 µm, amplitude ≈ 1 µm).
*Scale bars: (G, I, K, N) 100 µm. Measurements represent mean ± SD of at least three independent fields.*

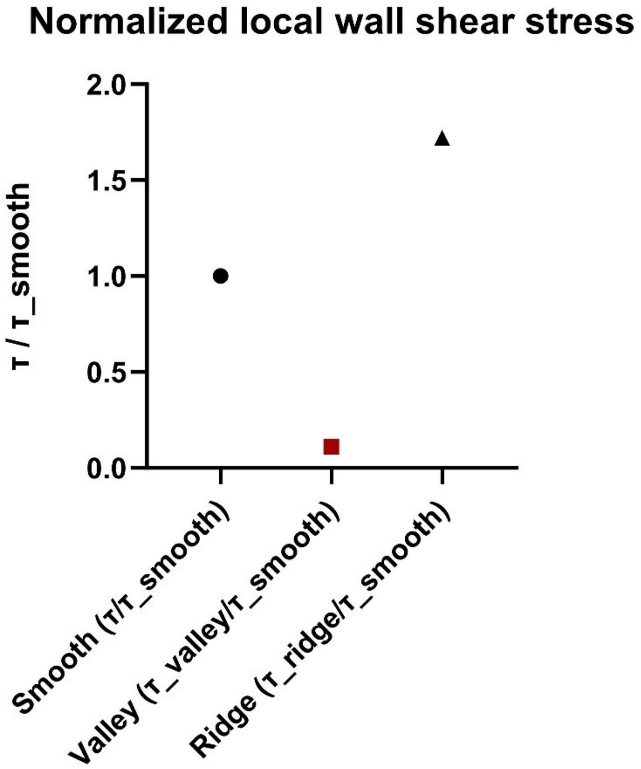


**Supplementary Figure S4 | Absolute local wall shear stress values in grooved and smooth lumens**
Absolute local wall shear stress (WSS, Pa) extracted from CFD simulations for smooth luminal regions and for groove valleys and ridge crests in the micro-grooved geometry. Values correspond to the same flow conditions as in Fig. 2 (10 ml·min$^{-1}$, 8 mm lumen diameter, Newtonian fluid). While the area-averaged wall shear stress remains matched between smooth and grooved lumens, local WSS is markedly reduced in groove valleys and elevated at ridge crests, giving rise to a broad local shear range.

**Supplementary Table S2 | Local wall shear stress values used for normalization**
Wall shear stress at the flat surface (τ1), inside the groove valley (τ2), and at the ridge crest (τ3) for all rheological models. Ratios τ2/τ1 and τ3/τ1 were used to compute normalized values presented in Fig. 2F and Supplementary Fig. S8.

| Model | $\tau_1$ [Pa] | $\tau_2$ [Pa] | $\tau_3$ [Pa] | $\tau_2/\tau_1$ | $\tau_3/\tau_1$ |
|---|---|---|---|---|---|
| M1 | 9.6 | 2.2 | 16.5 | 0.23 | 1.72 |
| M2 | 9.6 | 1.1 | 16.5 | 0.11 | 1.72 |
| M3 | 66.4 | 16.5 | 114.2 | 0.25 | 1.72 |

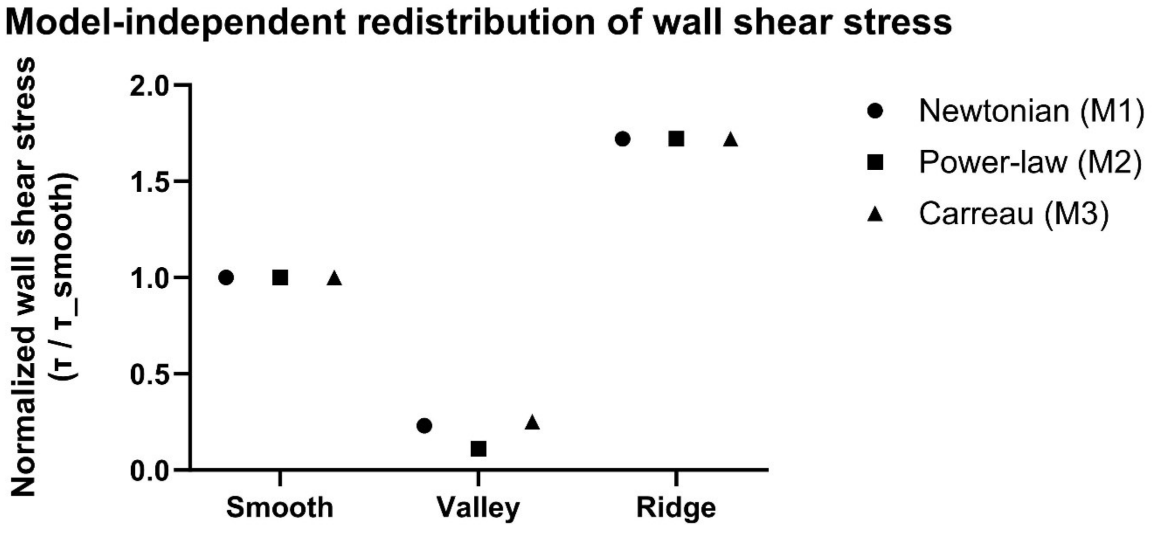


**Supplementary Figure S5 | Model-independent redistribution of wall shear stress by longitudinal micro-grooves**

Normalized local wall shear stress (τ / τ_smooth) for smooth wall regions, groove valleys, and ridge crests computed using three rheological descriptions of the perfusate: Newtonian (M1), power-law (M2), and Carreau (M3). Across all models, longitudinal micro-grooves consistently redistribute shear stress into low-shear valleys and high-shear ridges while preserving the mean wall shear stress, demonstrating that the effect is robust and independent of the specific rheological assumption.

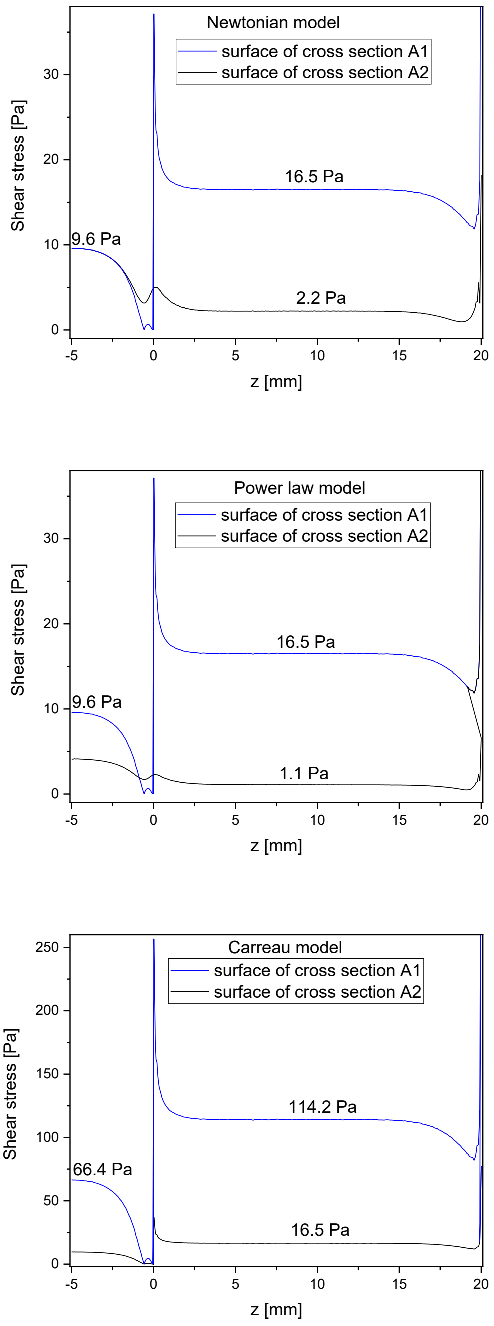


**Supplementary Figure S6 |** Wall shear stress profiles along cross-sections A1–A2
Wall shear stress (Pa) plotted along the longitudinal coordinate (z) for smooth and grooved luminal surfaces, extracted along cross-sections A1 (ridge-aligned) and A2 (groove valley) for three rheological models: Newtonian, power-law, and Carreau. Local minima correspond to groove valleys, whereas maxima correspond to ridge crests. Numerical values reported in Supplementary Table S2 were extracted from these profiles.

**Supplementary Note 1 | Rheological model definitions**
**Blood rheology was modeled using three constitutive descriptions:**

(i) Newtonian fluid with constant dynamic viscosity $\mu = 0.9$ mPa·s,
(ii) Power-law model defined as $\tau = K \cdot \dot{\gamma}^n$ ($n = 0.7$, $K = 0.016$ Pa·$s^n$),
(iii) Carreau model (parameters listed below).
Model comparison confirmed that, while absolute WSS values varied across rheological descriptions, the normalized redistribution pattern (valley-to-ridge ratio) remained consistent (Supplementary Fig. S7), demonstrating the robustness of the geometric effect.

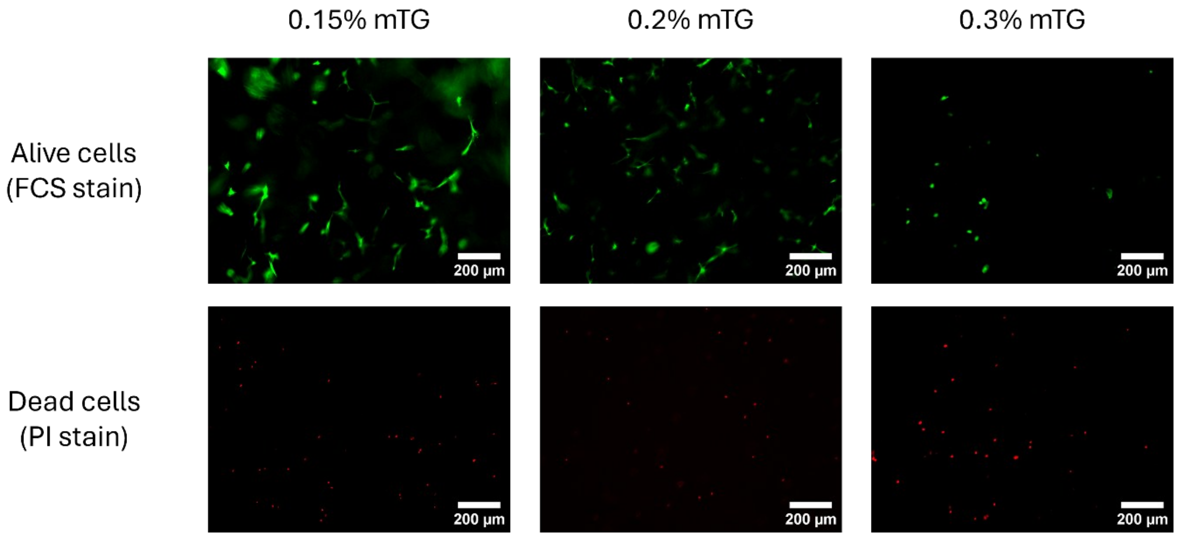


**Figure S7 | Concentration dependent mTG on cell shape and viability.**

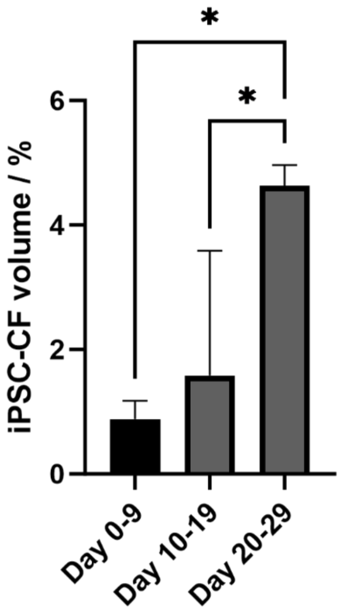


**Figure S8.** (D) Volume fraction analysis confirms progressive iPSC-CF tissue densification and cellular expansion from early seeding to mature constructs (mean ± SD, n = 6; one-way ANOVA + Tukey, $p < 0.05$).

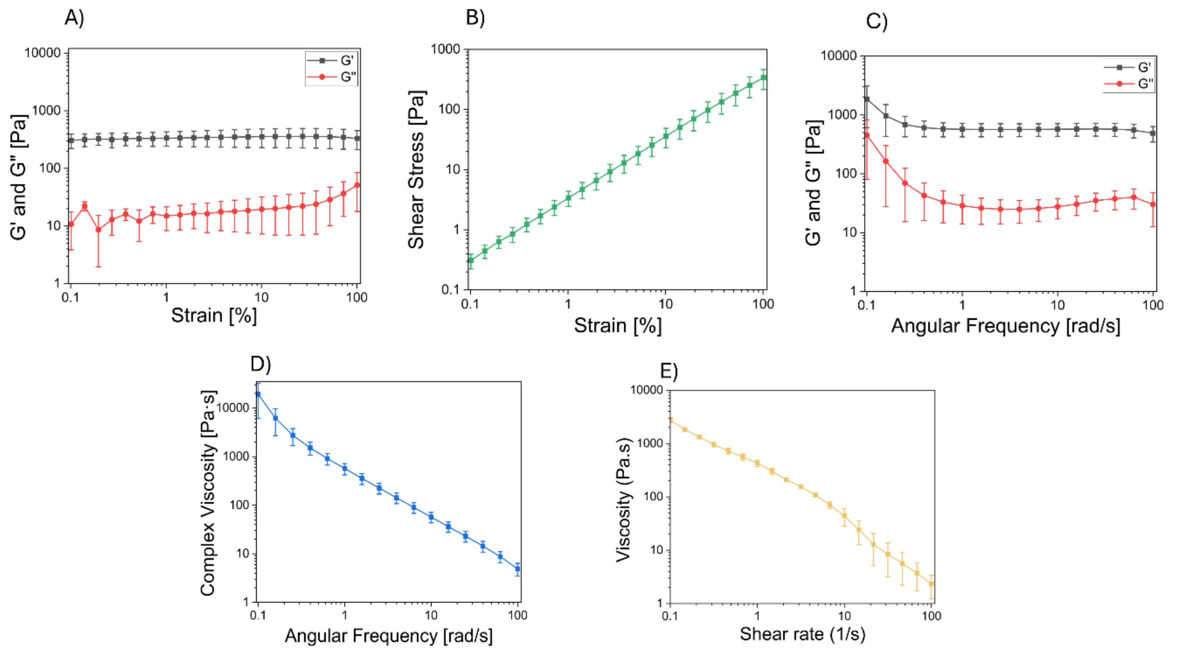


**Figure S9 | Rheological fingerprints and microstructure of ELR2-based composites.** (A) Amplitude sweep at 1 Hz showing dominant elastic behaviour (G′ > G″) and defining the linear viscoelastic region.
(B) Stress–strain curve illustrating nonlinear response and yield behaviour at high deformation.

(C) Frequency sweep confirming gel-like character with G′ > G″ across the entire range. (D) Complex viscosity versus angular frequency and (E) steady-shear viscosity versus shear rate demonstrate pronounced shear-thinning behaviour, favourable for extrusion-based printing.

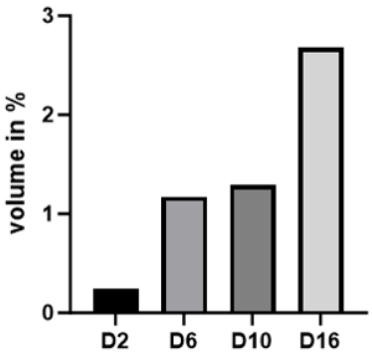


**Figure S10.** (D) Volume fraction analysis confirms progressive SMCs tissue densification and cellular expansion from early seeding to mature constructs (mean ± SD, n = 6; one-way ANOVA + Tukey, $p < 0.05$).

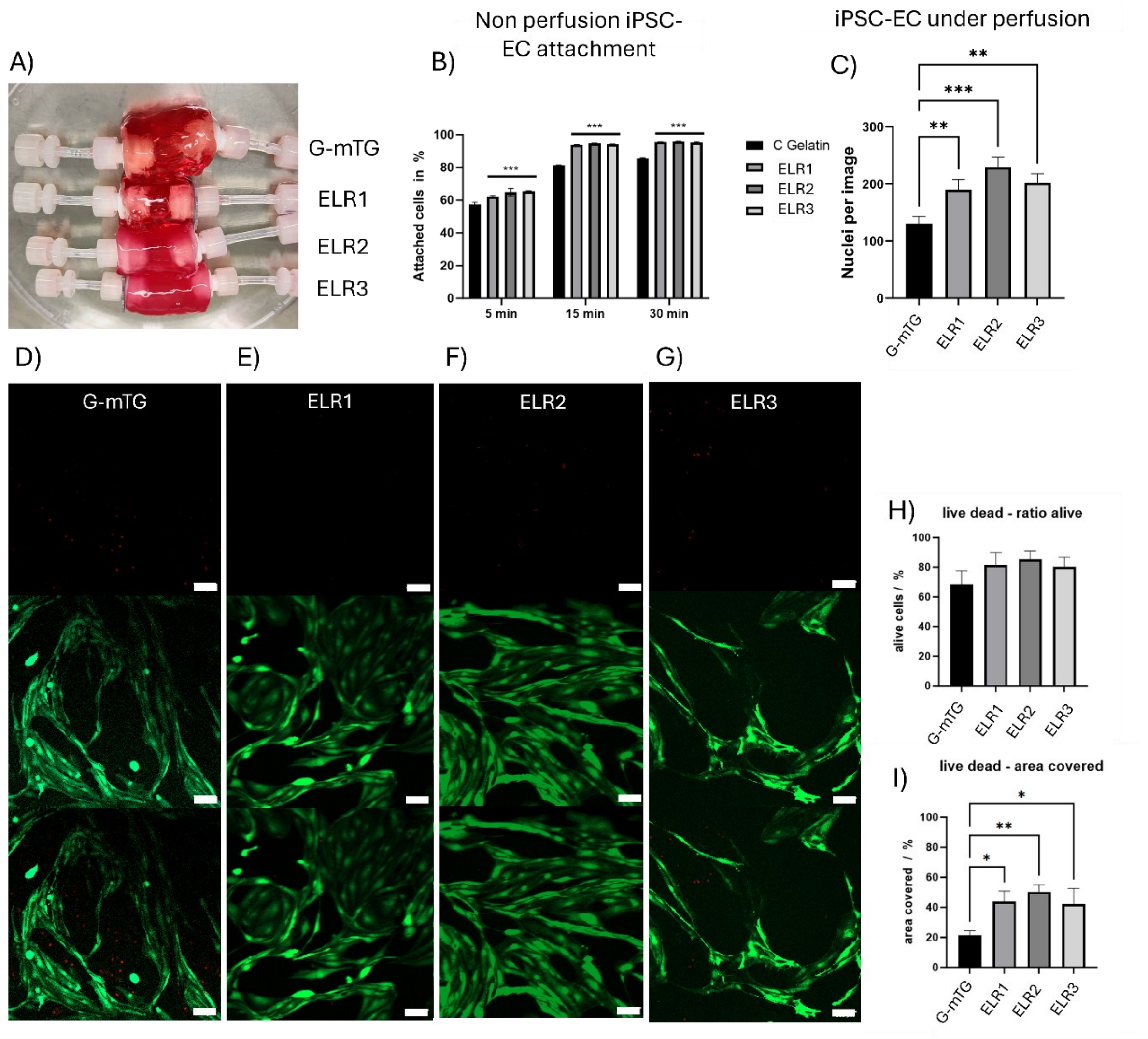


**Figure S11 | Pre-screening of smooth gelatin- and ELR-based hydrogels for early iPSC-EC attachment and viability.**

(A) Macroscopic view of triplicate tubular grafts (8 mm ID) composed of gelatin-mTG (G-mTG) and different ELR composites (ELR1, ELR2, ELR3).
(B) *Non-perfusion* attachment kinetics of iPSC-ECs over 5, 15 and 30 min on smooth hydrogels. ELSR-G supported the highest percentage of adherent cells at all timepoints compared with gelatin-mTG and other ELR blends (**** $p < 0.0001$, one-way ANOVA + Tukey).
(C) *Low-flow conditioning* of tubular grafts ($Q \approx 0.6$ mL min$^{-1}$; $\bar{\tau}w \approx 0.002$ dyn cm$^{-2}$) immediately after seeding, showing consistent cell attachment across triplicates. This represents a gentle circulation step used to prevent nutrient stratification and should not be interpreted as a retention assay (see Fig. 7 for 10 mL min$^{-1}$ shear).
(D–G) Representative Live/Dead confocal images after 24 h show > 90 % viability for all ELR groups and more extensive spreading on ELR (green = live, red = dead; scale = 100 µm).
(H–I) Quantification of (H) percentage of live cells and (I) surface coverage confirms significantly greater coverage for ELR2 than gelatin-mTG (** $p < 0.01$). These low-shear, smooth-surface screens identified ELR2 as the optimal bioink for endothelial seeding and formed the basis for patterned-lumen experiments under higher flow (10 mL min$^{-1}$) presented in Fig. 7.

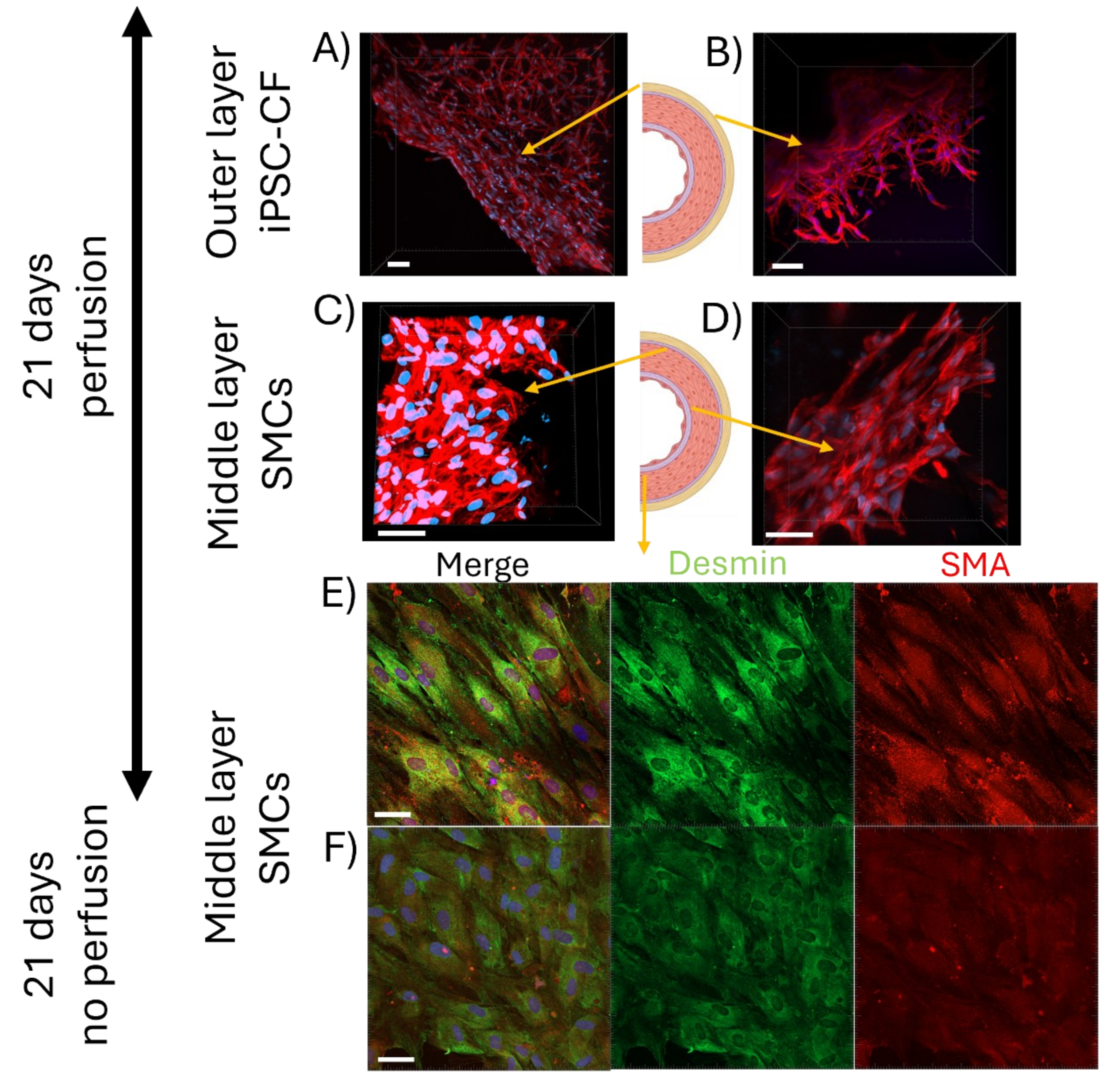


**Figure S12 | Layer-resolved organization and depth-dependent alignment of SMCs in tri-layer grafts after 21 days**

(A) Outer adventitial layer (iPSC-CFs) showing a dense, interwoven F-actin$^+$ network.
(B) Inner CF region spanning the adventitia–media interface, demonstrating structural continuity.
(C) Outer medial compartment (ELR2-based SMC layer) with interconnected α-SMA$^+$/desmin$^+$ bundles.
(D) Inner medial region adjacent to the lumen, where SMC bundles display preferential axial alignment parallel to the tube axis.
(E) En-face luminal view (construct halved longitudinally) under perfusion culture (21 days), revealing elongated, flow-aligned α-SMA$^+$/desmin$^+$ SMCs near the luminal interface.
(F) En-face control cultured without perfusion (21 days), showing shorter SMCs with largely random orientation.
Panels A–D: F-actin (red), nuclei (DAPI, blue).
Panels E–F: α-SMA (green), desmin (red), nuclei (DAPI, blue).
Scale bars = 50 µm.
